# Secondary metabolite profiling of *Pseudomonas aeruginosa* isolates reveals rare genomic traits

**DOI:** 10.1101/2023.12.18.572185

**Authors:** Rachel L. Neve, Emily Giedraitis, Madeline S. Akbari, Shirli Cohen, Vanessa V. Phelan

## Abstract

*Pseudomonas aeruginosa* is a ubiquitous gram-negative opportunistic pathogen with remarkable phylogenetic and phenotypic variability. In this work, we applied classical molecular networking analysis to secondary metabolite profiling data from seven *Pseudomonas aeruginosa* strains, including five clinical isolates from the lung secretions of people with cystic fibrosis. Combined with whole-genome sequencing, we show that some *P. aeruginosa* isolates, including nmFLRO1, produce a previously unreported class of acyl putrescines, isolate SH3A does not produce di-rhamnolipids because its genome belongs to phylogenetic clade 5, and the secondary metabolite profile of isolate SH1B reflects a frame-shift mutation in the quorum sensing regulator *rhlR.* This study highlights for the first time that secondary metabolite profiling provides unique insight into genetic variation of *P. aeruginosa*.

**Importance:** Secondary metabolite profiling of *Pseudomonas aeruginosa* isolates can be used to identify rare genomic variants that impact quorum sensing and metabolite biosynthesis that underlie virulence.

## Introduction

*Pseudomonas aeruginosa* is a ubiquitous gram-negative bacterium responsible for both acute and chronic infections. The remarkable phylogenetic and phenotypic diversity of *P. aeruginosa*, including varied virulence mechanisms, conversion to mucoid by overproduction of alginate, and altered quorum sensing (QS) pathways contribute to its survival in varied environments [1–5]. One mechanism *P. aeruginosa* uses to interact with its environment is through the production of secondary metabolites [6–8]. These compounds are known to mediate a variety of virulence mechanisms used by *P. aeruginosa* in the host environment, including swarming motility, QS, iron acquisition, and the production of reactive oxygen species [9–11]. Recently, liquid chromatography tandem mass spectrometry (LC-MS/MS) profiling of the *P. aeruginosa* secondary metabolome has been applied to identify markers of virulence, measure the effect of nutritional changes on QS pathways, and to capture the diversity of the metabolites produced [7–9, 12, 13].

The secondary metabolome of *P. aeruginosa* has been extensively characterized and consists of a small number of molecular families, including homoserine lactones (HSLs), phenazines (PHZs), alkyl quinolones (AQs), rhamnolipids (RLs), and the siderophores pyochelin (PCH) and pyoverdine (PVD) [7, 9, 11, 14]. The production of secondary metabolites by *P. aeruginosa* is dependent, in part, on the functionality of their biosynthetic gene clusters (BGCs) and the QS pathways that regulate their production [15]. The QS system of *P. aeruginosa* is composed of three interdependent regulatory circuits: *las, rhl,* and *pqs* systems [16]. Historically considered hierarchical, the three QS systems are inherently intertwined with overlapping regulons. In the *las* QS system, LasI produces the autoinducer 3-oxo-dodecanoyl-L-homoserine lactone (3-oxo-C12-HSL), which is subsequently bound by the LasR transcriptional regulator [17, 18]. LasR bound to 3-oxo-C12-HSL dimerizes and activates a variety of transcriptional pathways, including the *rhl* QS system. In *rhl* QS, RhlI produces *N*-butanoyl-L-homoserine lactone (C4-HSL), which binds to the RhlR transcriptional regulator, enabling dimerization and activation of its regulon [19]. The RhlI/RhlR system directly controls RL production [20]. Both *las* and *rhl* QS systems regulate the *pqs* QS system through the transcriptional regulator PqsR (MvfR) [16]. PqsR regulates the production of AQs, including the autoinducers 2-heptyl-4-quinolone (HHQ) and Pseudomonas quinolone signal (PQS), with PQS a more potent inducer of PqsR than HHQ. The BGC for the AQs includes the protein PqsE, which is not required for AQ production, but is required for production of the PHZs, including pyocyanin (PYO) [19]. The function of PqsE in QS has not been fully elucidated, but it is known to function as a chaperone providing stability to RhlR and may also produce an uncharacterized signaling molecule [21, 22]. Both RhlR and PqsE are required for production of PYO by *P. aeruginosa* [17, 18]. The BGCs of secondary metabolites are well conserved in the genomes of *P. aeruginosa* isolates, but mutations within the QS pathways are common. Mutations affecting LasR are more common than RhlR [23]. However, some isolates with LasR QS mutations are still capable of producing secondary metabolites by secondary mutations within other transcriptional regulators, including MexT, which can activate the *rhl* QS pathways [24].

Herein, we applied secondary metabolite profiling to five isolates of *P. aeruginosa* from cystic fibrosis (CF) sputum cultured in synthetic CF medium 2 (SCFM2), a medium developed to mimic the nutritional environment of the CF lung [25]. Approximately 60% of adults with CF have chronic and recurrent airway infections caused by *P. aeruginosa*, which contributes to morbidity and mortality [26]. By performing presence-absence analysis on the metabolomics data, we show that CF clinical isolates produce the same suite of secondary metabolites as laboratory strains PAO1 and PA14. However, some metabolites were only produced by one or more of the clinical isolates. Evaluating the metabolomics data for metabolites produced only by a single isolate of *P. aeruginosa*, we discovered that one of the isolates, mFLRO1, produces a class of acyl putrescines from the conjugation of putrescine with various fatty acids and another, SH3A, only produces mono-RLs due to its genome similarity to phylogenetic clade 5 representative strain PA-W1. By evaluating the metabolomics data for metabolites produced by all strains except one, we discovered that the genome of isolate SH1B has a one base frameshift mutation in *rhlR* that disrupts QS signaling, leading to decreased production of RLs and PHZs. Taken together, our data shows that profiling the secondary metabolome can be used to identify rare genomic variants that impact QS and metabolite biosynthesis in *P. aeruginosa* isolates.

## Results and Discussion

Five *P. aeruginosa* CF clinical isolates, including nmFLRO1, mFLRO1, SH1B, SH2D, and SH3A, were subjected to secondary metabolite profiling and whole genome sequencing (Table S1). Isolates nmFLRO1 and mFLRO1 were isolated at the University of California, San Diego Adult CF Clinic. Strains SH1B, SH2D, and SH3A were isolated from an individual with CF in Hanover, Germany before, during, and after a transient infection with *Burkholderia cenocepacia* H111, respectively [27]. The isolates were cultured in SCFM2 alongside laboratory strains PAO1 and PA14 and gross morphology of the cultures was visualized using a stereo microscope (Figure 1A). All *P. aeruginosa* strains formed macrostructures at the air-liquid interface of the cultures, which descended into aggregates on the bottom of the wells. Aligning with our previous results, PAO1 cultures in SCFM2 were blue in color, indicating production of the phenazine pyocyanin (PYO) [7]. The cultures of the other strains were less visually distinct, ranging in color from a moderate yellow (nmFLRO1) to faint green (SH3A) and colorless (mFLRO1). The color of the cultures and complexity of the macrostructures were strain-dependent and somewhat variable between replicates (Figure S1). Although most of the macro-aggregate structures were dispersed across the top of the well, two replicates of mFLRO1 formed a tighter pattern. The variability in culture color and aggregation between the strains was not due to differences in growth, as there was similar recovery of colony forming units (CFU/mL) after disruption of the aggregates from all cultures (Figure 1B). Although all cultures became alkaline, those of mFLRO1 were less basic than the other strains (Figure 1C).

**Figure 1.**
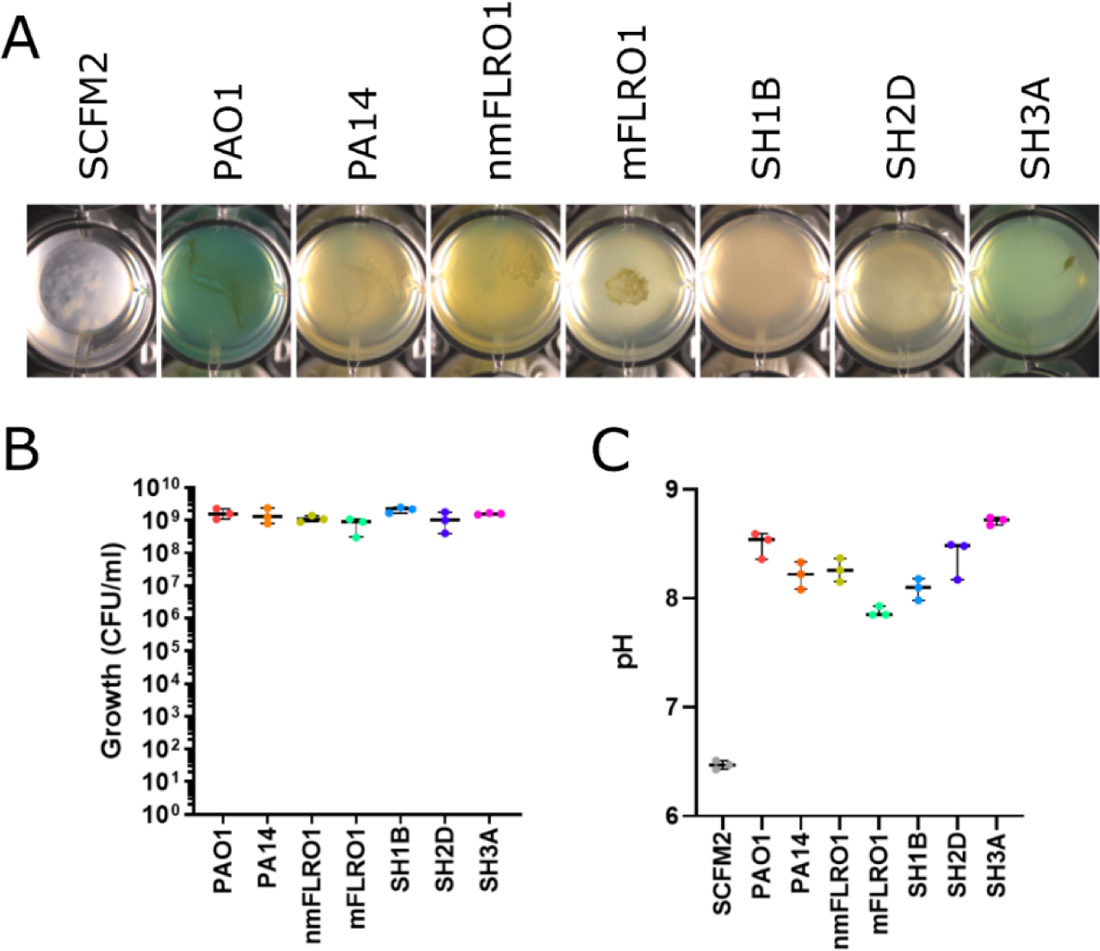
Growth of seven *P. aeruginosa* strains in SCFM2. (A) Representative phenotypes of strains grown statically in 1 mL of SCFM2 (*n* = 3) in a 48-well plate for 48 hr at 37 °C. Photographs of replicate growth wells are in Fig. S1 in the supplemental material. (B) Quantified growth of each strain as CFU/mL. (C) pH after mechanical disruption of cultures.

### Classical molecular networking analysis of isolates cultured in SCFM2

To determine whether the CF isolates produced the same suite of secondary metabolites as the laboratory strains, culture extracts were subjected to LC-MS/MS metabolomics and classical molecular networking (CMN) [28]. CMN organizes structurally related metabolites into molecular families by leveraging the concept that molecules with similar chemical structures have similar fragmentation patterns in their MS/MS spectra. In the molecular network, the MS/MS spectra of metabolites are represented as nodes and the relatedness between the MS/MS spectra is illustrated by the thickness of the connecting edges. Each node in a classical molecular network is most often composed of MS/MS spectra from multiple samples. Metadata groups denoting strain name (“Strain,” e.g., “PAO1”) and type (“Origin,” e.g., “Lab”) were used to identify which samples contributed to each node via MS/MS spectra. The number of nodes shared by each combination of sample groups was quantified and visualized as UpSet plots.

Four hundred nine molecular ions were captured in the classical molecular network (Figure 2A). Under the experimental conditions, four members of the PHZ molecular family were measured, including PYO, 1-hydroxyphenazine (1-HP), phenazine-1-carboxamide (PCN), and phenazine-1-carboxylic acid (PCA). Forty-two AQs were detected, including HHQ, 2-heptyl-4-hydroxyquinoline *N*-oxide (HQNO), PQS, and their associated nonyl congeners (NHQ, NQNO, and C9-PQS). RLs produced by *P. aeruginosa* consist of two sub-families, mono- and di-RLs with one and two rhamnose units bound to 3LJ(3LJhydroxyalkanoyloxy) alkanoic acids (HAAs), respectively. Both mono- and di-RLs were detected from various cultures, as was the siderophore PCH. The HSLs and PVD were below the limit of detection.

**Figure 2.**
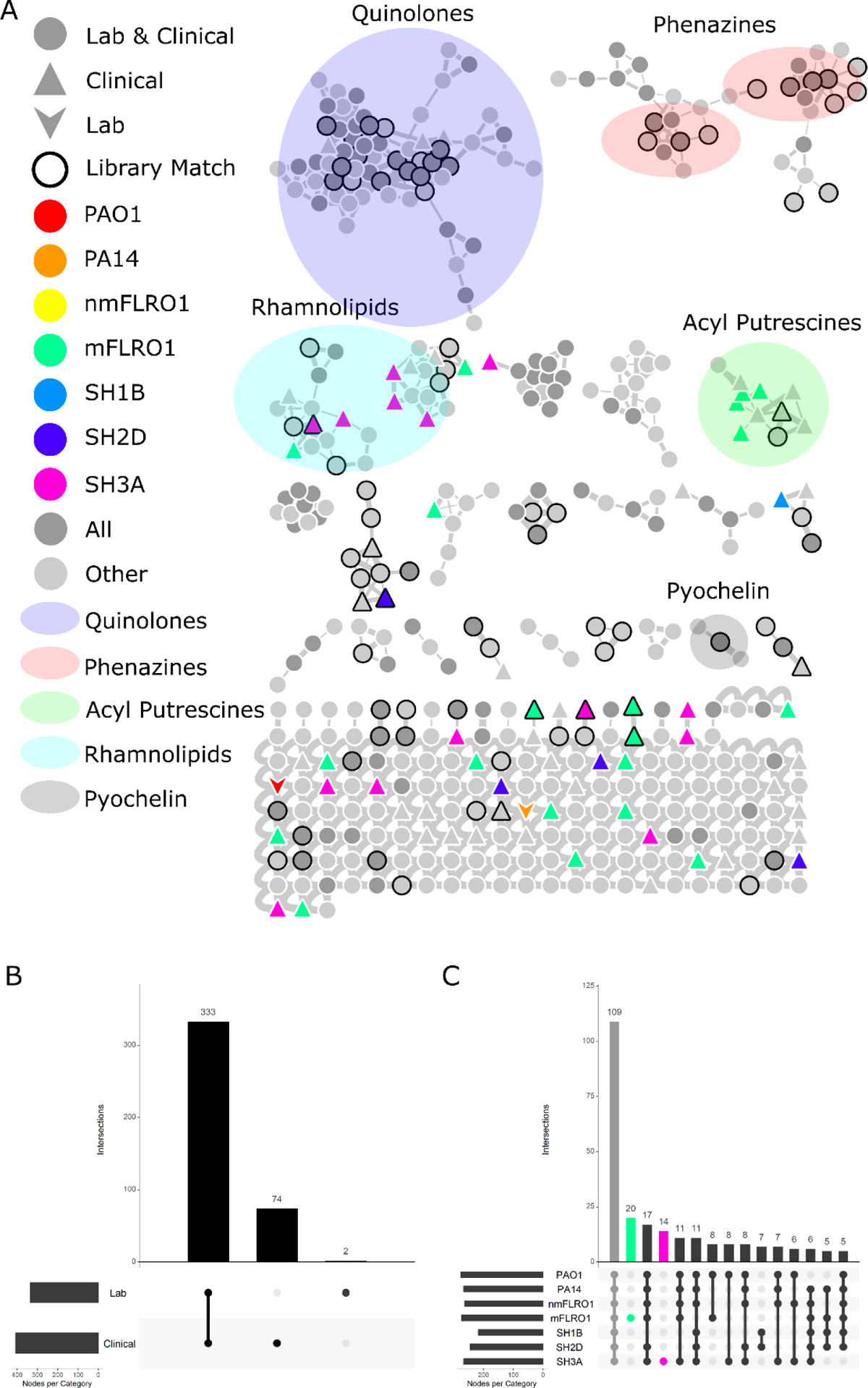
Molecular ions detected from the cultures of seven *P. aeruginosa* strains. (A) Classical molecular network of seven *P. aeruginosa* strains cultured in SCFM2. Nodes represent molecular ions (precursor mass and MS/MS spectra). The node shape represents whether the molecular ion was detected from cultures of laboratory strains (down arrow), clinical isolates (triangle), or both (circle). The node color indicates whether the molecular ion was detected in cultures from all strains (dark gray), individual strains (various colors), or combinations of strains (light gray). The widths of the lines connecting nodes (edges) represent the similarity of the MS/MS fragmentation of the connected nodes. Black node outlines indicate a spectral match between the data and MS/MS spectra within the GNPS spectral libraries. Nodes corresponding to the five molecular families discussed in the main text are highlighted by ovals. (B) UpSet plot quantifying the number of molecular ions detected from cultures of laboratory strains (PAO1 and PA14) compared to clinical isolates (nmFLRO1, mFLRO1, SH1B, SH2D, and SH3A). (B) UpSet plot quantifying the number of molecular ions detected from different combinations of cultures.

Three hundred thirty-three molecular ions (81%) were detected from the cultures of at least one laboratory strain and one clinical isolate (Figure 2B), with 109 nodes (26%) measured from the cultures of all strains (Figure 2C). Secondary metabolites detected from all strains included 1-HP, PCN, PCA, HHQ, HQNO, PQS, NHQ, NQNO, C9-PQS, and PCH. Notably absent from this set of secondary metabolites were the RLs and PYO. Seventeen nodes were detected in all strains except SH1B, including those representing RLs and PYO (Figure 2C). Although most nodes in the network represented shared molecular ions between lab and clinical strain cultures, 74 nodes (18%) were detected solely from the cultures of clinical isolates (Figure 2B), primarily from the cultures of mFLRO1 (20 nodes) and SH3A (14 nodes) (Figure 2C). Only two nodes were measured solely from the cultures of the laboratory strains (Figure 2B). This data suggests that *P. aeruginosa* isolates produce the same suite of secondary metabolites as laboratory strains; however, some clinical isolates produce unique compounds and/or are unable to produce specific secondary metabolites under these culture conditions.

### Clinical isolate mFLRO1 produces acyl putrescines

Twenty nodes in the classical molecular network represented molecular ions uniquely detected from the cultures of mFLRO1 (Figure 2). Four of these nodes clustered within a molecular family of 11 members (Figure 3A). Two nodes of the molecular family had spectral matches to the MS/MS spectra of acyl putrescine synthetic standards deposited within the GNPS libraries (Figure S2) [29]. Acyl putrescines consist of the polyamine putrescine conjugated to fatty acids of varying chain length and unsaturation. The two library matches putatively identified putrescine conjugated to hexadecenoic acid (putrescine C16:1) and octadecenoic acid (putrescine C18:1). These annotations were manually confirmed by comparing the experimental accurate mass and MS/MS measurements to the predicted and library values. The other nine members of the acyl putrescine molecular family were putatively annotated based on mass defect and MS/MS spectral similarity to the GNPS library matches.

**Figure 3.**
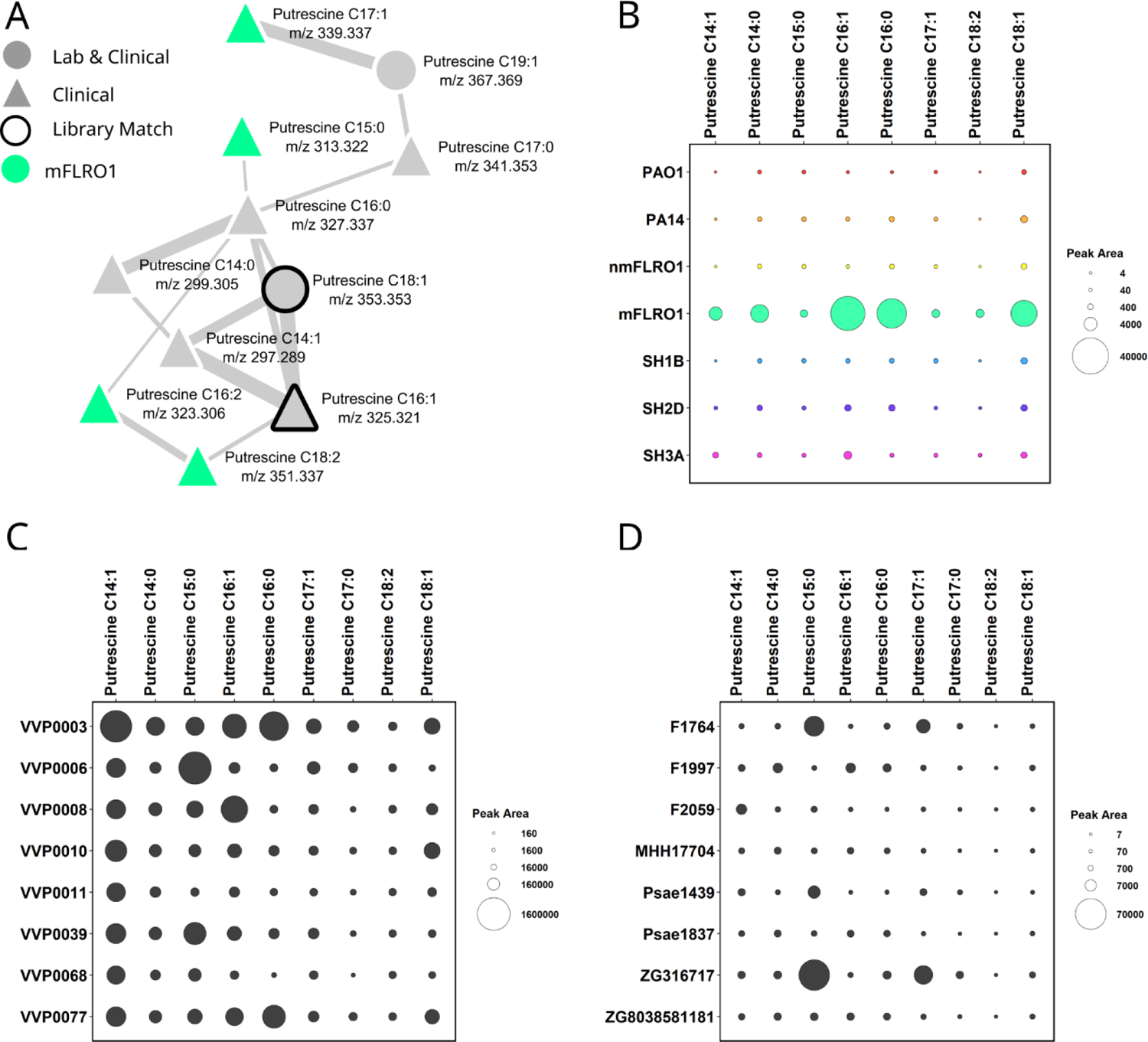
Acyl putrescine levels produced by *P. aeruginosa* isolates. (A) Acyl putrescin molecular family labeled with acyl chain length, unsaturation, and mass to charge ratio. Nodes represent molecular ions (precursor mass and MS/MS spectra). The node shape represent whether the molecular ion was detected from cultures of clinical isolates (triangle) or both laboratory strains and clinical isolates (circle). The node color indicates whether the molecular ion was detected in cultures from mFLRO1 (green) or combinations of strains (light gray). The widths of the lines connecting nodes (edges) represent the similarity of the MS/MS fragmentation of the connected nodes. Black node outlines indicate a spectral match between the data and MS/MS spectra within the GNPS spectral libraries. (B) Bubble plot representing the average quantitation (peak area) across biological replicates (*n* = 3) from each *P. aeruginosa* strain for the eight acyl putrescines above the limit of quantitation. Corresponding box plots for each metabolite are in Figure S3. (C) Bubble plot representing the quantitation (peak area) of nine acyl putrescines from Dataset 1 (MSV000086107): publicly available LC-MS/MS data capturing the secondary metabolome of eight *P. aeruginosa* strains isolated from CF sputum cultured in LB under standard laboratory conditions (*n* = 1 biological replicate). (D) Bubble plot representing the quantitation (peak area) of nine acyl putrescines from eight *P. aeruginosa* strains from Dataset 2 (MSV000089869): publicly available LC-MS/MS data capturing the secondary metabolome of 35 *P. aeruginosa* isolates selected from the Helmholtz Centre for Infection Research biobank which were isolated from various sites of infection and cultured in LB under standard laboratory conditions (*n* = 1 biological replicate). Acyl putrescine levels from all 35 isolates are displayed in Figure S4.

Of the eleven nodes within the acyl putrescine molecular family, nine nodes were composed of spectra solely from the cultures of clinical isolates. To determine which of the isolates produced the highest quantity of acyl putrescines, their relative abundance was quantified and compared between strains (Figure 3B, Figure S3). Only eight of the acyl putrescines were measured above the limit of quantitation. Strain mFLRO1 produced the highest concentration of acyl putrescines, preferentially producing putrescine C16:1, putrescine C18:1, and putrescine conjugated to hexadecanoic acid (putrescine C16:0).

To determine whether other *P. aeruginosa* isolates could produce acyl putrescines, two publicly available metabolomics datasets were downloaded from the MassIVE data repository and queried using MassQL. Datasets MSV000086107 and MSV000089869 contain LC-MS/MS data that captured the secondary metabolome of eight *P. aeruginosa* clinical isolates collected at the University of California, San Diego Adult CF Clinic and 35 isolates from various sites of infection selected from the Helmholtz Centre for Infection Research biobank, respectively [9, 13]. The data for both datasets were collected from *P. aeruginosa* isolates cultured in LB.

MassQL is a computational method that enables users to search publicly available metabolomics data for patterns captured in MS data that are intrinsically related to compound structure, such as isotopic ratios or compound class-specific fragments captured in the MS/MS spectra [30]. The MS/MS spectra of datasets MSV000086107 and MSV000089869 were searched for the presence of two structural characteristics of acyl putrescines captured in their MS/MS spectra: a neutral loss corresponding to the mass of NH_2_ (−17.0264 Da) and a fragment ion of m/z 72.0802 corresponding to the molecular formula C_4_H_12_N^+^. This MassQL query resulted in the identification of acyl putrescines in both datasets. These results were manually verified by comparing experimental accurate mass and MS/MS measurements between the queried dataset and the annotations of the acyl putrescines described above.

The acyl putrescines were quantified from all samples within each dataset using the built-in quantitation algorithm of GNPS Dashboard [31]. GNPS Dashboard is a web-based platform for the visualization of open-source format LC-MS data. Chromatographic peak area for individual molecules is calculated in GNPS Dashboard using user provided exact mass and retention time parameters. Importantly, due to differences in experimental conditions and data acquisition, intensity values could only be compared within each dataset. Statistical comparison of acyl putrescine abundance between the isolates included in the datasets could not be performed as both datasets contained metabolomics data for single replicates.

Acyl putrescines were detected from all eight clinical isolates within dataset MSV000086107, with VVP0003 and VVP0006 producing the highest relative quantities (Figure 3C). Unlike mFLRO1, which predominantly produced putrescine C16:1, putrescine C18:1, and putrescine C16:0, the fatty acids incorporated into the acyl putrescines by the isolates of dataset MSV000086107 varied in length and unsaturation. Isolate VVP0003 preferentially produced putrescine C14:0, putrescine 16:1, and putrescine 16:0; VVP0006 produced putrescine C15:0, and VVP0008 produced putrescine C16:1. Acyl putrescines were also detected in culture extracts of Dataset MSV000089869 isolates (Figure 3D, Figure S4). Of the 35 isolates included in the study, strains F1764 (isolated from the respiratory tract) and ZG316717 (isolated from an ear infection) produced the highest levels of acyl putrescines, with both preferentially making putrescine C15:0 and putrescine C17:1. This result showed that different *P. aeruginosa* isolates produce acyl putrescines, with strain-dependent incorporation of different fatty acids.

Dysregulation of polyamine biosynthesis in *P. aeruginosa* clinical isolates is well documented [32–36]. Putrescine is catabolized to succinate and ammonia through the putrescine utilization pathway and/or converted to spermidine by *P. aeruginosa*. To control free spermidine levels, *P. aeruginosa* is known to *N-*acetylate spermidine [37, 38]. However, neither acetylated nor acylated spermidine was detected in mFLRO1 culture extracts. Production of acyl putrescines by *P. aeruginosa* isolates may be an alternative deactivation mechanism to control culture alkalinity due to putrescine degradation, as evidenced by the lower alkalinity of mFLRO1 cultures compared to the other isolates (Figure 1C), and/or to reduce the toxicity associated with high free spermidine levels [39, 40].

### Clinical isolate SH3A is genetically related to PA-W1

Fourteen nodes in the classical molecular network represented molecular ions uniquely detected from the cultures of SH3A (Figure 2). Three of these nodes clustered within a molecular family of 13 members (Figure 4A). Three nodes within the molecular family had spectral matches to the MS/MS spectra of characterized mono- and di-RLs RC10C10, RRC10C10, and RRC10C12:1 in the GNPS libraries. The other ten members of the RL molecular family were annotated based on mass defect and MS/MS spectral similarity to the GNPS library matches. This analysis led to the identification of seven mono-RLs and three di-RLs, including the discovery of one mono-RL and one di-RL with monoacetylated rhamnose units (Figure S5).

**Figure 4.**
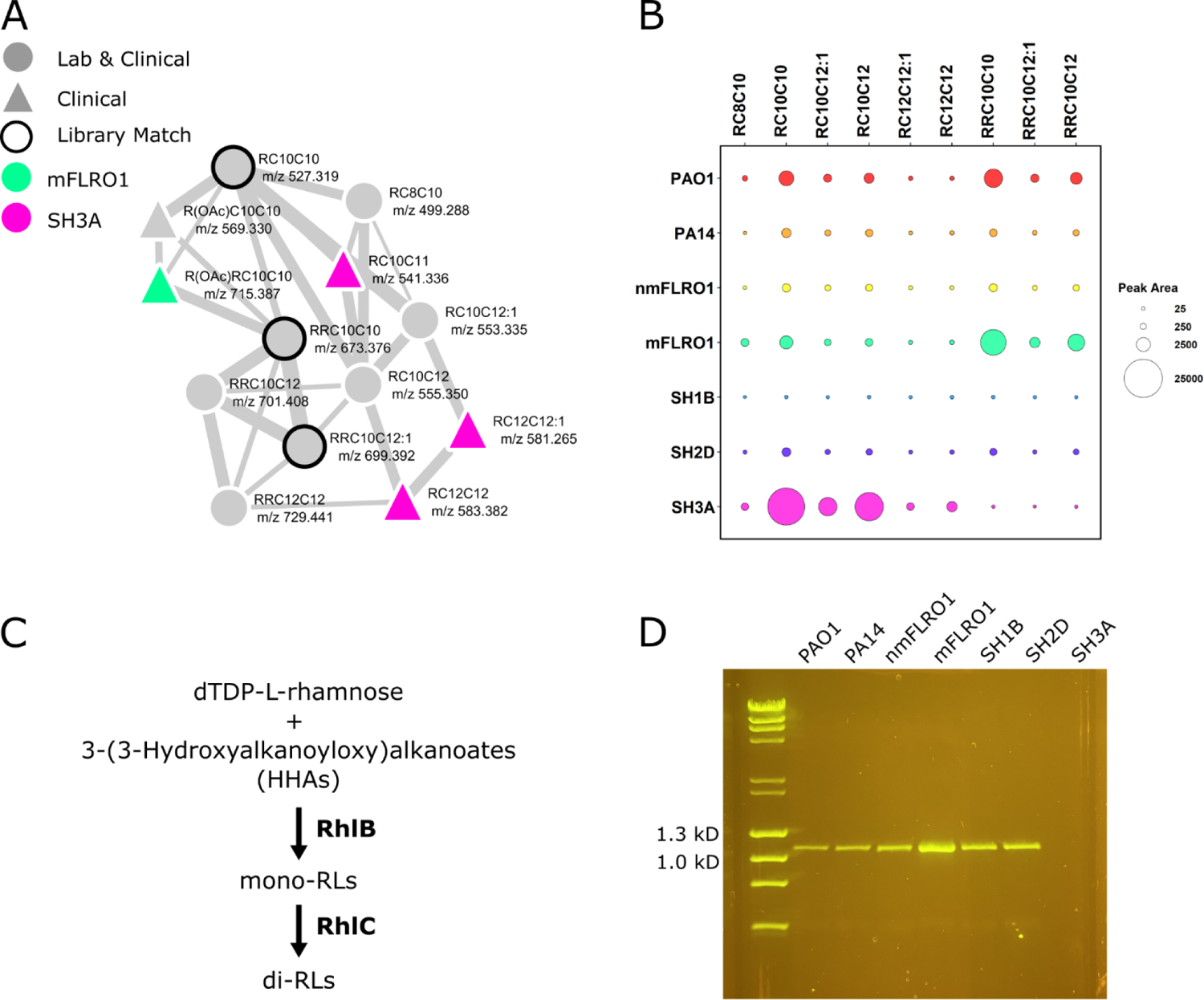
Rhamnolipid production by *P. aeruginosa* strains. (A) Rhamnolipid molecular family labeled with one or two rhamnose units (R or RR, respectively), acyl chain length, unsaturation, and mass to charge ratio. Nodes represent molecular ions (precursor mass and MS/MS spectra). The node shape represents whether the molecular ion was detected from cultures of clinical isolates (triangle) or both laboratory strains and clinical isolates (circle). The node color indicates whether the molecular ion was detected uniquely in cultures from SH3A (pink) or mFLRO1 (green), or combinations of strains (gray). The widths of the lines connecting nodes (edges) represent the similarity of the MS/MS fragmentation of the connected nodes. Black node outlines indicate a spectral match between the data and MS/MS spectra within the GNPS spectral libraries. (B) Bubble plot representing the average quantitation (peak area) of biological replicates (*n* = 3) from each *P. aeruginosa* strain for the nine rhamnolipids above the limit of quantitation. Corresponding box plots for each metabolite are in Figure S6. (C) Biosynthetic pathway of mono-RLs and di-RLs from precursors. (D) PCR analysis for the presence of *rhlC* in the genomes of the seven *P. aeruginosa* strains.

The three nodes representing molecular ions in the RL molecular family detected from only SH3A cultures were annotated as three mono-RLs (RC10C11, RC12C12:1, and RC12C12). Further evaluation of the metadata information in category “Strain” of the molecular family revealed that although SH3A produced mono-RLs, di-RLs were not detected from SH3A cultures. To determine whether preferential production of mono-RLs was unique to SH3A, RL relative abundance was quantified and compared between strains (Figure 4B, Figure S6). Only nine of the RLs were measured above the limit of quantitation. Strain SH3A was the only strain to exclusively produce mono-RLs.

In *P. aeruginosa*, di-RLs are produced by the sequential functions of RhlA, RhlB, and RhlC (Figure 4C). RhlA produces HAAs, which are subsequently glycosylated by the rhamnosyltransferase RhlB to form mono-RLs [41]. RhlC condenses dTDP-L-rhamnose with mono-RLs to produce di-RLs. Although RhlA and RhlB are encoded in a bicistronic operon (*rhlAB*), RhlC is encoded separately on the *P. aeruginosa* genome [42]. The inability of SH3A to produce di-RLs led to the hypothesis that SH3A did not have functional RhlC, resulting in the production of only mono-RLs. PCR analysis of the SH3A genome revealed that it did not encode for RhlC (Figure 4D).

Previously reported phylogenetic analysis of thousands of *P. aeruginosa* strains showed that the genomes of isolates segregate into five distinct phylogenetic clades, where the genomes of PAO1, PA14, PA7, and PA-W1 are representative examples of clades 1, 2, 3, and 5, respectively [43–45]. Isolates belonging to phylogenetic clades 3-5 represent less than 1% of the sequenced genomes of *P. aeruginosa* and are genetically distinct from laboratory strains PAO1 and PA14, with some researchers advocating for these isolates to be considered a separate species [46]. Despite compelling evidence of the existence of clade 4, it has not yet been independently confirmed [43].

To determine if the genome of SH3A belonged to clade 3 or 5, the presence or absence of the differentiating genes was evaluated between the genomes of PAO1 (clade 1), PA14 (clade 2), PA7 (clade 3), PA-W1 (clade 5) and SH3A (Table 1). The genomes of clades 1, 2, 3, and 5 can be distinguished from each other by the presence or absence of a subset of genes, including *exoS, exoY, phzH,* genes for the type III secretion system (T3SS), *rhlC, exlA, exlB, catA, oprA,* and *fosA* [44, 47]. Unlike clade 1 and 2 genomes, clade 3 and 5 genomes do not contain genes for *exoS*, *exoY, phzH, rhlC,* and the T3SS, but have genes for *exlA, exlB* and *oprA* [43, 45]. Evaluation of the presence of these genes revealed that the SH3A genome is a member of clade 3 or clade 5 of the *P. aeruginosa* phylogenetic tree.

**Table 1.** Genes distinguishing SH3A as a Clade 5 isolate.

|  | PAO1<br>Clade 1 | PA14<br>Clade 2 | PA7<br>Clade 3 | PA-W1<br>Clade 5 | SH3A |
| --- | --- | --- | --- | --- | --- |
| ***exoS*** | + | - | - | - | - |
| ***exoY*** | + | + | - | - | - |
| ***phzH*** | + | + | - | - | - |
| ***T3SS*** | + | + | - | - | - |
| ***rhlC*** | + | + | - | - | - |
| ***exlAB*** | - | - | + | + | + |
| ***oprA*** | - | - | + | + | + |
| ***catA*** | - | - | + | - | - |
| ***fosA*** | + | + | - | + | + |
| ***PA1132*** | + | + | - | + | + |
| ***PA1133*** | + | + | - | + | + |

Clade 3 and 5 genomes can be differentiated from each other by the presence of *catA* in clade 3 genomes but absent from clade 5 genomes and *fosA* absent in clade 3 genomes, but present in clade 5 genomes. Notably, the deletion of *rhlC* from the clade 3 and 5 genomes are thought to have occurred through different evolutionary events [44]. In clade 3 genomes, the loss of *rhlC* is the result of a five gene deletion event, including *fosA, rhlC, PA1131, PA1132,* and *PA1133*. In clade 5 genomes, the loss of *rhlC* is thought to result from a smaller deletion event, leaving *fosA*, *PA1132,* and *PA1133* intact. This analysis revealed that SH3A has the genes *fosA, PA1132,* and *PA1133* intact; meaning it does not produce di-RLs because the SH3A genome belongs to clade 5 of the *P. aeruginosa* phylogenetic tree.

Like the genomes of PA7 and PA-W1, *rhlC* and *phzH* are absent from the genome of SH3A. RhlC is required for di-RL production and PhzH is required to produce the PHZ PCN [42, 48]. Similar to *rhlC*, *phzH* is encoded in the *P. aeruginosa* genome in a location distinct from the core phenazine biosynthetic pathways [48]. As the isolates representing clades 3 and 5 are poorly represented in publicly available genomic databases, screening the secondary metabolome of libraries of *P. aeruginosa* isolates for loss of di-RL and PCN production despite production of the mono-RLs and the other PHZs could be used to prioritize strains that putatively belong to the phylogenetic clades 3 and 5 for genome sequencing, enabling more thorough phylogenetic comparison between isolates within the clade.

### Clinical isolate SH1B genome has disrupted *rhlR*

Seventeen nodes in the classical molecular network represented molecular ions that were detected from all cultures, except those of SH1B (Figure 2). Three of these nodes clustered within the RL molecular family and were annotated as the mono-RLs RC10C10, RC10C12:1, and RC10C12. Further evaluation of the metadata information in category “Strain” of the molecular family revealed that no RLs were detected from SH1B cultures. Although SH1B produced AQs and PCH, the PHZ PYO was not detected from SH1B cultures. To verify this observation, the secondary metabolites 1-HP, PYO, PCA, HHQ, HQNO, PQS, RC10C10, RRC10C10, and PCH were quantified from SH1B cultures and compared to levels from PA14 (Figure 5A, Figure S7). Indeed, SH1B produced markedly reduced levels of PHZs, RLs, and PQS compared to PA14, but continued to produce HHQ, HQNO, and PCH. This pattern of secondary metabolite production was reminiscent of the results from the secondary metabolite profiling of an *rhlR* deletion mutant compared to its PA14 parental strain [49, 50].

**Figure 5.**
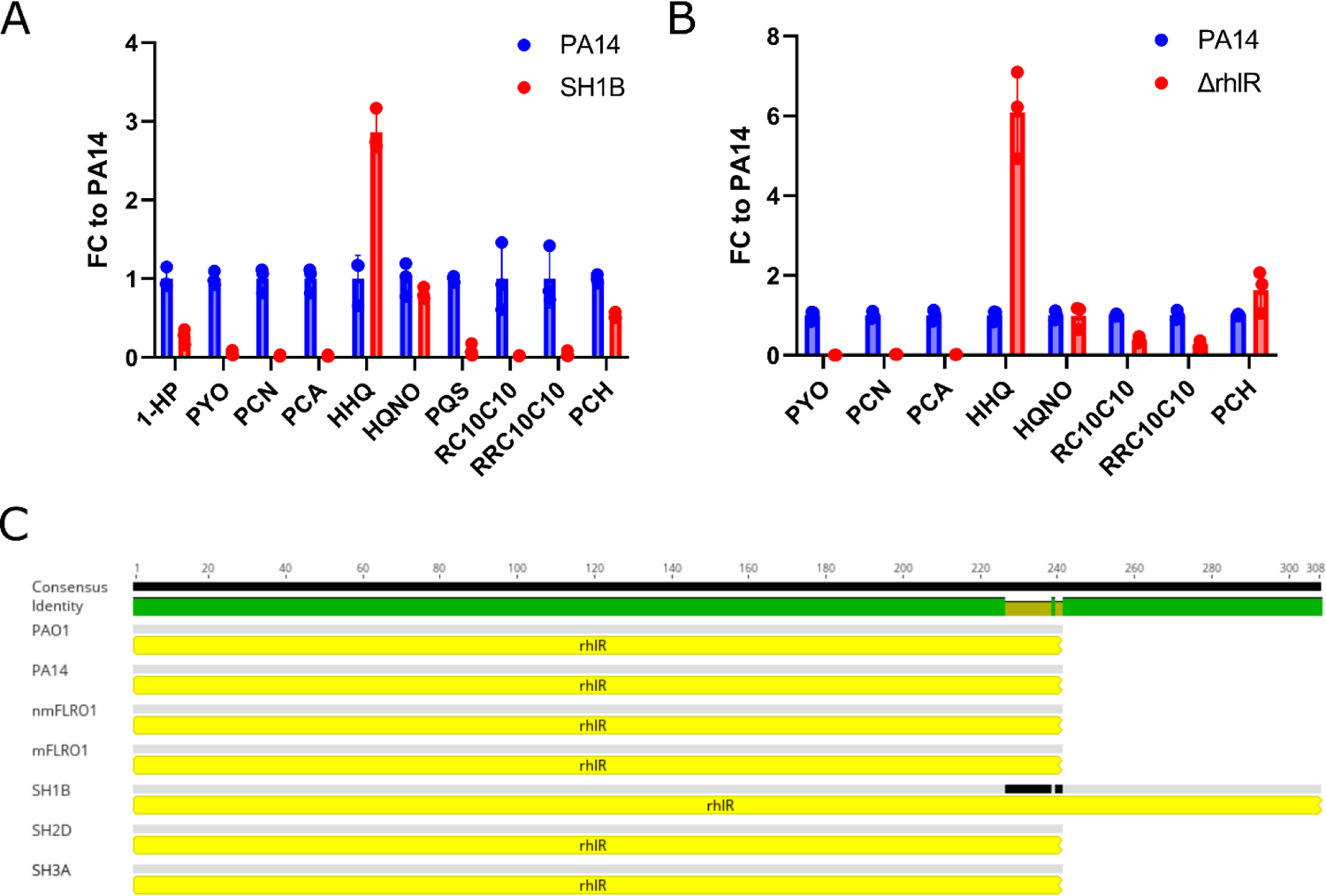
SH1B has disrupted *rhl* QS. (A) Ratio of secondary metabolite levels produced by SH1B compared to PA14 in SCFM2. Corresponding box plots for each metabolite are in Figure S7. (B) Ratio of secondary metabolite levels produced by Δ*rhlR* compared to the PA14 parental strain in LB (MSV000083500). 1-HP and PQS were below the limit of detection and/or quantitation. Corresponding box plots for each metabolite are in Figure S8. (C) The *rhlR* gene of SH1B is predicted to produce an elongated RhlR protein compared to PAO1, PA14, nmFLRO1, mFLRO1, SH2D, and SH3A due to a single nucleotide insertion, which leads to a one-base frame shift and improper termination of translation. Corresponding gene sequence alignment is in Figure S9.

To confirm that the pattern of SH1B secondary metabolite production reflected the pattern of the *rhlR* deletion mutant, the publicly available metabolomics dataset MSV000083500 was downloaded and processed. Dataset MSV000083500 consists of LC-MS/MS data that captured the secondary metabolome of Δ*rhlR* and its parental strain PA14 cultured in LB in biological triplicate [50]. Comparison of the secondary metabolite levels between Δ*rhlR* and wild-type PA14 (WT) indicated that although Δ*rhlR* produced reduced quantities of RLs and PHZs, it produced AQs and PCH at levels equivalent to or higher than WT (Figure 5B, Figure S8). The similarity of the secondary metabolite profile of SH1B to Δ*rhlR* led to the hypothesis that SH1B had disrupted *rhl* QS signaling.

To determine if the genome of *P. aeruginosa* isolate SH1B had a mutation in the *rhlR* gene, the nucleotide sequence of the SH1B *rhlR* was compared to the sequences from the genomes of the other strains in this study. This analysis revealed that the SH1B *rhlR* gene sequence contained a one-base insertion frame-shift mutation (678insC), which disrupts the stop codon and leads to improper termination of RhlR protein synthesis (Figure 5C, Figure S9). The SH1B *rhlR* is predicted to produce a protein consisting of 308 amino acids, which is 67 amino acids longer than the RhlR produced by most *P. aeruginosa* strains. Moreover, the sequence for the SH1B *rhlR* stop codon is within the nucleotide sequence of *rhlI*, fully abrogating *rhl* QS.

To evaluate the frequency of this mutation among *P. aeruginosa* isolates, the nucleotide and predicted amino acid sequences for SH1B *rhlR* were searched against the genome sequences of *P. aeruginosa* isolates deposited into NCBI and Pseudomonas Genome Database [51, 52]. This analysis identified four other isolates with elongated *rhlR* gene sequences, including PA103 (isolated from sputum), BL22 (isolated from human eye), PABL048 (isolated from blood), and PS1793 (isolated from sputum). The *rhlR* nucleotide sequences of SH1B, PA103, BL22, PABL048, and PS1793 are 100% identical (Figure S10). This analysis indicated that the *rhlR* frame-shift mutation identified in the SH1B genome is rare (<0.5%) in the sequenced genomes of *P. aeruginosa.* Taken together, the inability of SH1B to produce RLs and PYO, but maintain production of AQs and PCH is due to a loss of function mutation in the *rhl* QS signaling pathway.

These results suggest that secondary metabolite profiling of *P. aeruginosa* isolates could be used to screen isolate libraries for mutants with defective QS pathways. Mutations within the QS signaling pathways of *P. aeruginosa* frequently occur, with mutations in *lasR* more common than mutations in *rhlR* [49]. The *las* QS system controls a regulon of virulence-associated genes, including the biosynthesis of all secondary metabolite molecular families [3]. Therefore, *P. aeruginosa* isolates with true loss of *lasR* function would be expected to be unable to produce any secondary metabolites. Despite the frequency of *lasR* loss of function mutations, some isolates with these mutations are still able to produce QS-regulated virulence factors [53]. In evolution experiments of laboratory strain *P. aeruginosa* PAO1, mutations within the gene of virulence-associated transcriptional regulator MexT recovered RL and PYO biosynthesis in isolates with a loss of function *lasR* mutation, providing an explanation for the origin of LasR-independent *rhl* QS pathway activation [24]. As a result, *P. aeruginosa* isolates with predicted *lasR* and/or *rhlR* loss of function mutations could be screened for the absence of the production of specific secondary metabolite classes, enabling the separation of true loss of QS function isolates from those that have escape mutations in *mexT* or other regulatory genes and provide additional insight into the association of genotypes with virulence.

## Conclusion

In this work, we used untargeted LC-MS/MS to profile the secondary metabolome of seven *P. aeruginosa* strains, including five uncharacterized clinical isolates (nmFLRO1, mFLRO1, SH1B, SH2D, and SH3A) from the sputum of people with CF and two laboratory strains (PAO1 and PA14). By analyzing the data with CMN, we quantified the commonalities and differences in secondary metabolite production between *P. aeruginosa* isolates and laboratory strains. This analysis revealed that *P. aeruginosa* isolates and laboratory strains produce a common secondary metabolome, but the ability to produce the different secondary metabolite molecular families is dictated, in part, by the genome sequence of the strain under study. We identified that some *P. aeruginosa* isolates, including mFLRO1, can produce a class of acyl putrescines and posit the production of this molecular family is a mechanism to deactivate putrescine degradation. Genomic sequencing of SH3A illustrated that its lack of di-RL production was indicative of its genome belonging to clade 5 of the *P. aeruginosa* phylogenetic tree. Using secondary metabolite profile similarity to Δ*rhlR*, we discovered that the genome of SH1B has a loss of function mutation in RhlR that disrupts *rhl* QS signaling, resulting in loss of RL and PYO biosynthesis. These results indicate that secondary metabolite profiling of *P. aeruginosa* isolates can provide unique insight into their genomic variability. Considering that this study profiled the secondary metabolome of five *P. aeruginosa* clinical isolates and captured three rare genetic traits representing <1-5% of the population with publicly available genomes or metabolomes, it raises the question of how truly infrequent these traits are in the population and whether the isolates captured in the current databases accurately represent the genomic diversity of *P. aeruginosa*. Future investigations into *P. aeruginosa* population diversity should integrate secondary metabolite profiling to aid in the prioritization of rare genotypes for sequencing, discern the effect of mutations on QS pathways, and be integrated with phenotypic screening to identify biomarkers of virulence.

## Materials and Methods

### Strains

The *P. aeruginosa* strains used in this study are summarized in Table S1.

### Culture of *P. aeruginosa* in SCFM2

Synthetic Cystic Fibrosis Medium 2 (SCFM2) was prepared as previously described with the following alteration: commercial bovine submaxillary mucin type I-S (BSM, Millipore Sigma) was dialyzed prior to addition to the medium [25].

Briefly, BSM was suspended in 1X 3-morpholinopropane-1-sulfonic acid (MOPS) buffer (pH 7.0), dialyzed against 1X MOPS using a 10 kD molecular weight cut-off cassette (Thermo Fisher Scientific), and sterilized using a liquid autoclave cycle (15-minute sterilization time) prior to addition to the medium. SCFM2 was stored in the dark at 4°C, checked for sterility prior to use, and used within one month of preparation. *P. aeruginosa* was inoculated from streak plates into 5 mL LB broth and incubated overnight at 37°C, shaking at 220 RPM. 0.990 µL of SCFM2 were inoculated with 10 µL of *P. aeruginosa* LB culture (OD600 of 0.05; ∼1×10^6^ CFU/mL) in a polystyrene 48 well plate. The plate was covered with its lid and incubated statically, under ambient oxygen conditions, at 37°C for 48 hrs.

### Sample Processing

Gross phenotypes of SCFM2 cultures were photographed with a top-down view of each well using a Stemi 508 stereo microscope with an Axiocam 105 color camera (Zeiss). Samples were mechanically disrupted by pipetting. Disrupted samples were aliquoted for growth measurements (CFU/mL), culture pH, and metabolomics analysis. All biological replicates (n = 3) were serially diluted, spotted onto LB agar, incubated at 37°C overnight, and counted to determine colony forming units by volume (CFU/mL).

### Sample Preparation for Metabolomics Analysis

Each sample was chemically disrupted with an equal volume of 1:1 solution of ethyl acetate (EtOAc, VWR HiPerSolv Chromanorm) and methanol (MeOH, Fisher Scientific Optima LC/MS grade). The samples were dried and stored at −20°C until use. After thawing, samples were resuspended in 100% MeOH, diluted 5-fold in 100% MeOH containing 1 µM glycocholic acid (Calbiochem, 100.1% pure), and centrifuged for 10 min at 4000 RPM (Thermo Sorvall ST 40R) to remove non-soluble particulates prior to injection.

### LC-MS/MS Data Acquisition

Mass spectrometry data acquisition was performed using a Bruker Daltonics Maxis II HD qTOF mass spectrometer equipped with a standard electrospray ionization (ESI) source as previously described [7]. The mass spectrometer was tuned by infusion of Tuning Mix ESI-TOF (Agilent Technologies) at a 3 µL/min flow rate. For accurate mass measurements, a wick saturated with Hexakis (1H,1H,2H-difluoroethoxy) phosphazene ions (Apollo Scientific, *m/z* 622.1978) located within the source was used as a lock mass internal calibrant. Samples were introduced by an Agilent 1290 UPLC using a 10 µL injection volume.

Extracts were separated using a Phenomenex Kinetex 2.6 µm C18 column (2.1 mm x 50 mm) using a 9-minute, linear water-acetonitrile (ACN) gradient (from 98:2 to 2:98 water:ACN) containing 0.1% formic acid at a flow rate of 0.5 mL/min. The mass spectrometer was operated in data dependent positive ion mode, automatically switching between full scan MS and MS/MS acquisitions. Full scan MS spectra (*m/z* 50 - 1500) were acquired in the TOF and the top five most intense ions in a particular scan were fragmented via collision induced dissociation (CID) using the stepping function in the collision cell. LC-MS/MS data for PA mix, a mixture of available commercial standards of *P. aeruginosa* secondary metabolites, were acquired under identical conditions. Bruker Daltonics CompassXport was used to apply lock mass calibration and convert the LC-MS/MS data from .d format to .mzXML format.

### Classical Molecular Networking (CMN)

The Molecular Networking workflow (version release 26) was applied to the .mzXML files using the GNPS analysis platform [28]. Briefly, the data were filtered by removing all MS/MS fragment ions ± 17 Da of the precursor m/z. MS/MS spectra were window filtered by choosing only the top 6 fragment ions in each ± 50 Da window throughout the spectrum. The precursor ion mass tolerance was set to 0.05 Da and the MS/MS fragment ion tolerance to 0.1 Da. Each node was set to consist of at least 5 matching spectra with a minimum of 5 matched fragment ions between nodes to generate an edge. A network was then created where edges between nodes were filtered to have a cosine score above 0.8. Further, edges between two nodes were kept in the network only if each of the nodes appeared in each other’s respective top 10 most similar nodes. Finally, the maximum size of a molecular family was set to 80, and the lowest scoring edges were removed until each molecular family was below this threshold. The spectra in the network were then searched against the GNPS spectral libraries.

Library spectra were filtered in the same manner as the input data. All matches kept between the network spectra and library spectra were required to have a cosine score above 0.7 and at least 5 matched peaks.

### UpSet Plot Generation

The network table generated via CMN was downloaded from Cytoscape. A binary presence/absence table was created from the network table to determine whether each node was detected in the cultures of each strain according to the GNPSGROUP designations. The binary table was visualized in R using the UpSetR package [22, 23].

### Secondary Metabolites Quantitation from CF Isolates

MZmine (version 2.53) was used to perform feature finding on the .mzXML files [54]. The output files included a feature table containing area under the curve (AUC) integration values for each feature (*m/z*-RT pair) and an .mgf file containing MS/MS fragment ions for each feature. Features were normalized by row sum and the mean aggregation was applied to peak abundances for each group. Features were filtered for rhamnolipids, acyl putrescines, and *P. aeruginosa* secondary metabolites using exact mass and retention time.

### Secondary metabolite analysis of PA14 and Δ*rhlR*

The publicly available *P. aeruginosa* secondary metabolite profiling dataset MSV000083500 was downloaded from MassIVE [50, 55]. MZmine (version 2.53) was used to perform feature finding on the .mzML files [54]. The output files included a feature table containing area under the curve (AUC) integration values for each feature (*m/z*-RT pair) and an .mgf file containing MS/MS fragment ions for each feature. Features were filtered for known *P. aeruginosa* metabolites.

### Metabolite Annotation

The classical molecular network (https://gnps.ucsd.edu/ProteoSAFe/status.jsp?task=de0b3f2b01174831b5a2be8546812ca1) was visualized using Cytoscape (version 3.7.1) [56]. Extracted ion chromatograms of m/z values of interest were visually inspected using the GNPS Dashboard (https://dashboard.gnps2.org/) and MS/MS spectra were visualized using the Metabolomics Spectrum Resolver (https://metabolomics-usi.gnps2.org/) [31, 57]. All annotated nodes corresponding to *P. aeruginosa* specialized metabolites identified from CMN are listed in Table S2. Annotations of metabolites corresponding to commercial standards (level 1 annotation) were confirmed by comparing the experimental data (exact mass, MS/MS, and retention time) with data acquired for the compound in PA mix using Bruker Daltonics DataAnalysis v4.1 (Build 362.7) [58]. Putative annotation of metabolites of interest corresponding to matches to the GNPS libraries (level 2 annotation) were validated by comparing the experimental data (exact mass, MS/MS) to reported data and putative structures, respectively [28, 58–60].

### MassQL Query of Public Metabolomics Datasets for Acyl Putrescines

Two published, publicly available *P. aeruginosa* secondary metabolite profiling datasets, MSV000086107 and MSV000089869, were downloaded from MassIVE to a GNPS workspace [9, 13, 28, 55]. These datasets, along with our own dataset as a positive control, were queried using MassQL v31.4 with the query: QUERY scaninfo(MS2DATA) WHERE MS2PROD=72.0802:TOLERANCEMZ=0.005 AND MS2NL=17.0264:TOLERANCEMZ=0.005 [31]. The results of the MassQL (v31.4) query (https://gnps.ucsd.edu/ProteoSAFe/status.jsp?task=23c630b0fc484582a5e0187c6ce5ef99) were subsequently networked using the Classical Molecular Networking workflow (v30) within GNPS (https://gnps.ucsd.edu/ProteoSAFe/status.jsp?task=f4284d9fa3f14ef6bceb6cbad0580033). The results were filtered using library matches to synthetic acyl putrescine standards in the GNPS libraries and precursor m/z values of the manually annotated acyl putrescines. To identify which isolates produced acyl putrescines, the chromatographic peak for each of the identified putrescines was identified in individual files using GNPS Dashboard and the putative annotations were verified by manual inspection of the associated MS/MS spectra [31, 57]. Peak areas of the acyl putrescines were quantified across each dataset using the built-in quantitation algorithm of GNPS Dashboard with the parameters of exact mass of the eight acyl putrescine with a 0.05 Da window and a retention time window of 30 seconds spanning peak maxima. The quantified values were visualized as bubble plots using R.

### PCR of *16S rRNA* and *rhlC* Genes

Genomic DNA was extracted from *P. aeruginosa* strains using the QIAamp DNA Mini Kit (Qiagen) according to manufacturers’ instructions. PCR-based detection of the *16S rRNA* (positive control) and *rhlC* genes was completed using the Phusion High-Fidelity PCR kit (ThermoFisher) following the manufacturer’s instructions with published primer sequences [42, 61]. Gel electrophoresis was performed on all samples using a 1.0% agarose gel in 1x TAE buffer and visualized using SYBR Safe DNA Gel Stain (ThermoFisher).

### Genome Sequencing

*P. aeruginosa* CF isolates were inoculated from streak plates into 5 mL LB broth and incubated overnight at 37°C, shaking at 220 RPM. Genomic DNA was extracted using the QIAamp DNA Mini Kit (Qiagen) according to manufacturers’ instructions and concentration and quality were confirmed by NanoDrop spectroscopy. Genomic sequencing using short-read Illumina and long-read Oxford Nanopore technologies (ONT), de novo assembly, and annotation was performed by SeqCenter (Pittsburgh, PA). Briefly, Porechop (v0.2.4) was used to trim residual adapter sequence from the ONT reads using default parameters that may have been missed during base calling and demultiplexing [62]. De novo genome assemblies were generated from the ONT read data with Flye (v2.9.2) under the nano-hq (ONT high-quality reads) model (--asm-coverage 50 --genome-size 6000000) [63]. Additional Flye options initiate the assembly by first using reads longer than an estimated N50 based on a genome size of 6 Mbp. Subsequent polishing used the Illumina read data with Pilon (v1.24) under default parameters [64]. To reduce erroneous assembly artifacts caused by low quality nanopore reads, long read contigs with an average short read coverage of 15x or less were removed from the assembly. Assembled contigs were evaluated for circularization via Circulator (v1.5.5) using the ONT long reads [65]. Assembly annotation was then performed with Bakta (v1.8.1) using the Bakta v5 database [66]. Assembly statistics were recorded with QUAST (v5.2.0) [67].

### Genomic Comparisons

Assembled and annotated genomes of the CF isolates included in this study were visualized in Geneious v2023.2.1 [68]. To determine if the genome of SH3A belonged to clade 3 or 5 of the *P. aeruginosa* phylogenetic tree, genomes of PAO1, PA14, PA7, and PAO1 were downloaded from NCBI and used as reference genomes for clades 1, 2, 3, and 5, respectively [51]. Full genomic alignments were conducted using the Mauve plug-in with default parameters [69]. Annotations of aligned genomes were searched for *exoS*, *exo*Y, *phzH*, *rhlC*, *exlA, exlB, oprA, catA, fosA, PA1131, PA1132, and PA1133* and the gene cluster for the type 3 secretion system (T3SS). To determine if *rhlR* of SH1B contained a mutation that would disrupt *rhl* QS signaling, the nucleotide and predicted protein sequences were extracted from the genomes of PAO1, PA14, nmFLRO1, mFLRO1, SH1B, SH2D, and SH3A and aligned using Clustal Omega v1.2.2 with default parameters [70]. The nucleotide and predicted protein sequences of SH1B *rhlR* were subsequently used for a Blast search in NCBI and the Pseudomonas Genome Database to identify other isolates with the same mutation [51, 52]. The nucleotide sequences of *rhlR* from isolates BL22, PA103, PABL048, and PS1793 were downloaded. The nucleotide and predicted protein sequences of *rhlR* from these four isolates were aligned to *rhlR* from SH1B and PAO1 using Clustal Omega v1.2.2 with default parameters [70].

### Statistical Analysis

Statistical comparison of metabolite abundance was conducted in GraphPad Prism (version 9.3.1). Unpaired two-tailed T-tests of differential secondary metabolite abundance between PA14 and SH1B and PA14 and Δ*rhlR* are summarized in the supplemental figure legends. For all analyses, P values of <0.05 were considered statistically significant.

## Data Availability

Mass spectrometry data files are available via MassIVE as MSV000087157. Genome sequences for nmFLRO1, mFLRO1, SH1B, SH2D, and SH3A are available in NCBI under BioProject XXXXXX (in process).

