## Supplemental Data for "Secondary metabolite profiling of *Pseudomonas aeruginosa* isolates reveals rare genomic traits"

Supplementary Material for

“Classical molecular networking of the secondary metabolome of *Pseudomonas aeruginosa* isolates reveals rare genomic traits”

Rachel L. Neve^1,*^, Emily Giedraitis^2,*^, Madeline Akbari^1^, Shirli Cohen^1^, Vanessa V. Phelan^2#^

**AUTHOR AFFILIATIONS:**

^1^ Department of Immunology and Microbiology, School of Medicine, University of Colorado - Anschutz Medical Campus, Aurora, CO, 80045, USA

^2^ Department of Pharmaceutical Sciences, Skaggs School of Pharmacy and Pharmaceutical Sciences, University of Colorado - Anschutz Medical Campus, Aurora, CO, 80045, USA

*Authors contributed equally to the study

**CORRESPONDING AUTHOR:**

[Figure S4 Bubble plot of acyl putrescine abundance of 35 strains from MSV000089869](#_Toc148003780)	6

[Table S2 Secondary metabolite annotation	1](#_Toc148003786)3

### Table S1 *P. aeruginosa* strains used in this study.

| **Strain** | **Origin** | **Other names** | **Source** |
| --- | --- | --- | --- |
| PAO1 | Lab strain | MPAO1 | University of Washington, Seattle |
| PA14 | Lab strain |  | Suzanne Noble,  University of California, San Francisco |
| nmFLRO1 | CF isolate |  | Forest Rohwer,  San Diego State University |
| mFLRO1 | CF isolate |  | Forest Rohwer,  San Diego State University |
| SH1B | CF isolate | Sh34, 10/92 | Leo Eberl,  ETH Zürich |
| SH2D | CF isolate | Sh37, 8/93 | Leo Eberl,  ETH Zürich |
| SH3A | CF isolate | Sh46, 9/95 | Leo Eberl,  ETH Zürich |

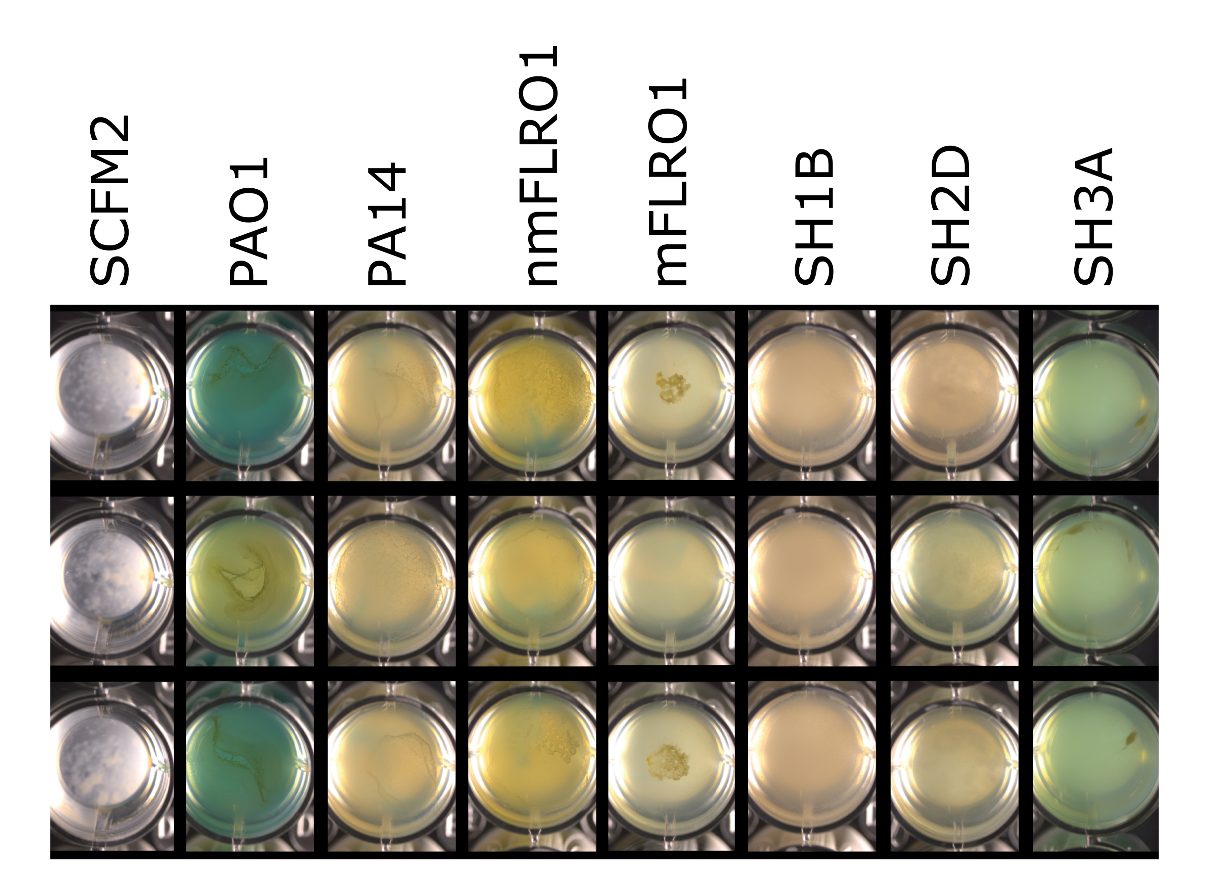

**Figure S1.** Growth of P. aeruginosa strains in SCFM2. Photographs of replicate control media wells and growth in SCFM2.

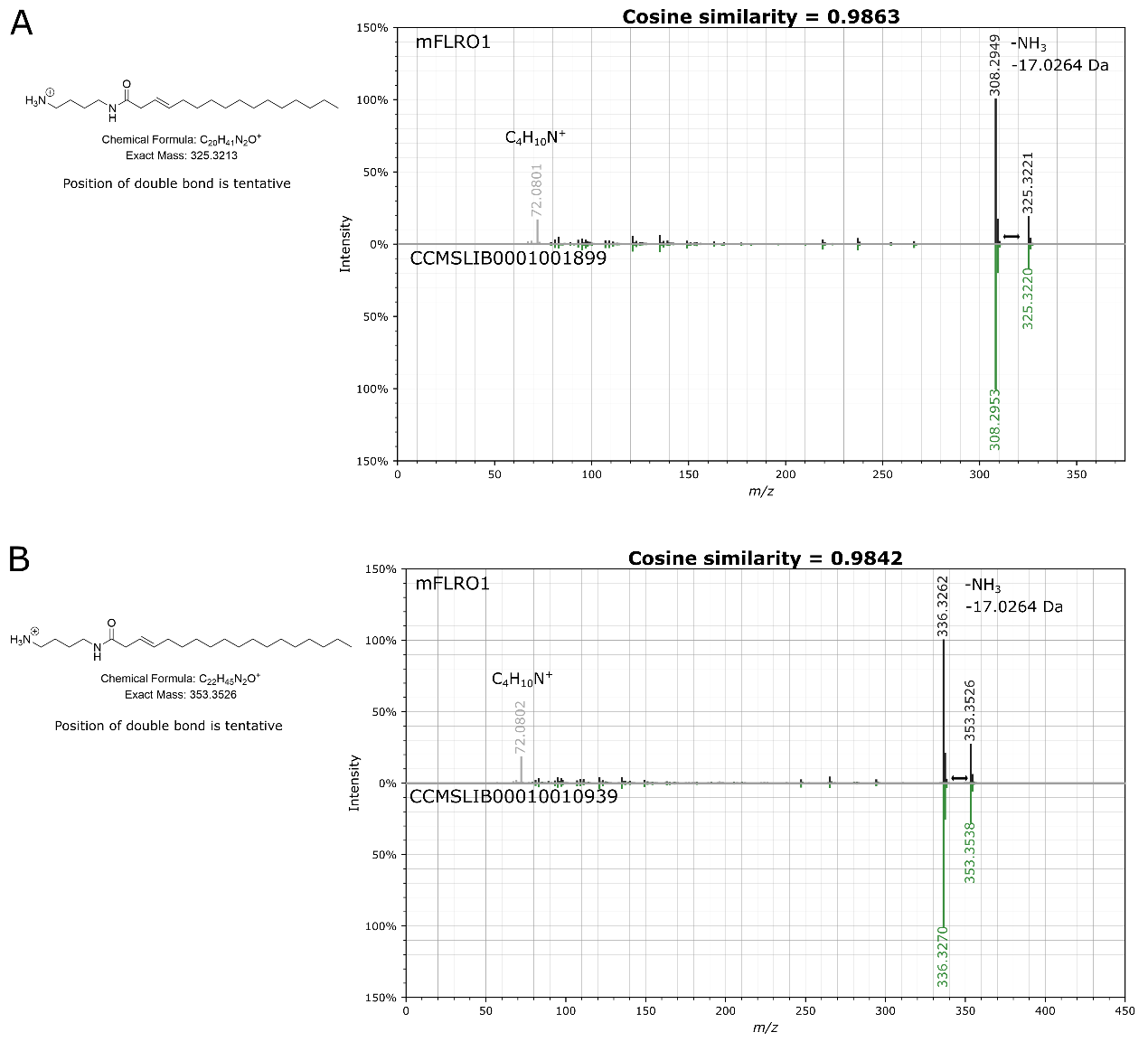

Figure S2. Comparison of MS/MS spectra between data collected from mFLRO1 cultures and GNPS spectral libraries for Putrescine C16:1 (A) and Putrescine C18:1 (B).

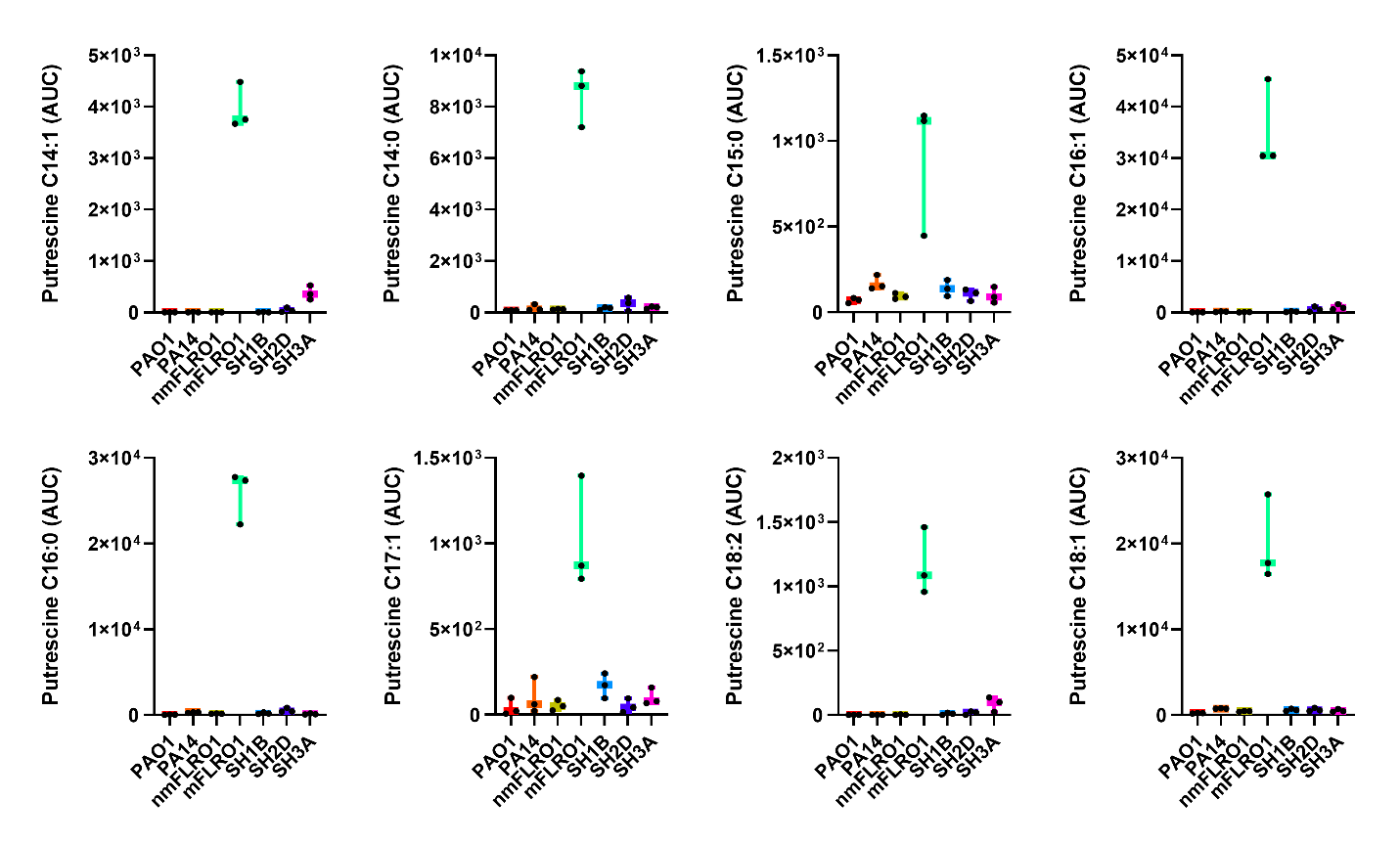

Figure S3. Differential levels of acyl putrescine congeners produced by seven P. aeruginosa strains in SCFM2. Box plots represent the 25 to 75th percentiles, with a line at the median. Error bars indicate the minimum to maximum. Individual sample values shown (n = 3 biological replicates per strain).

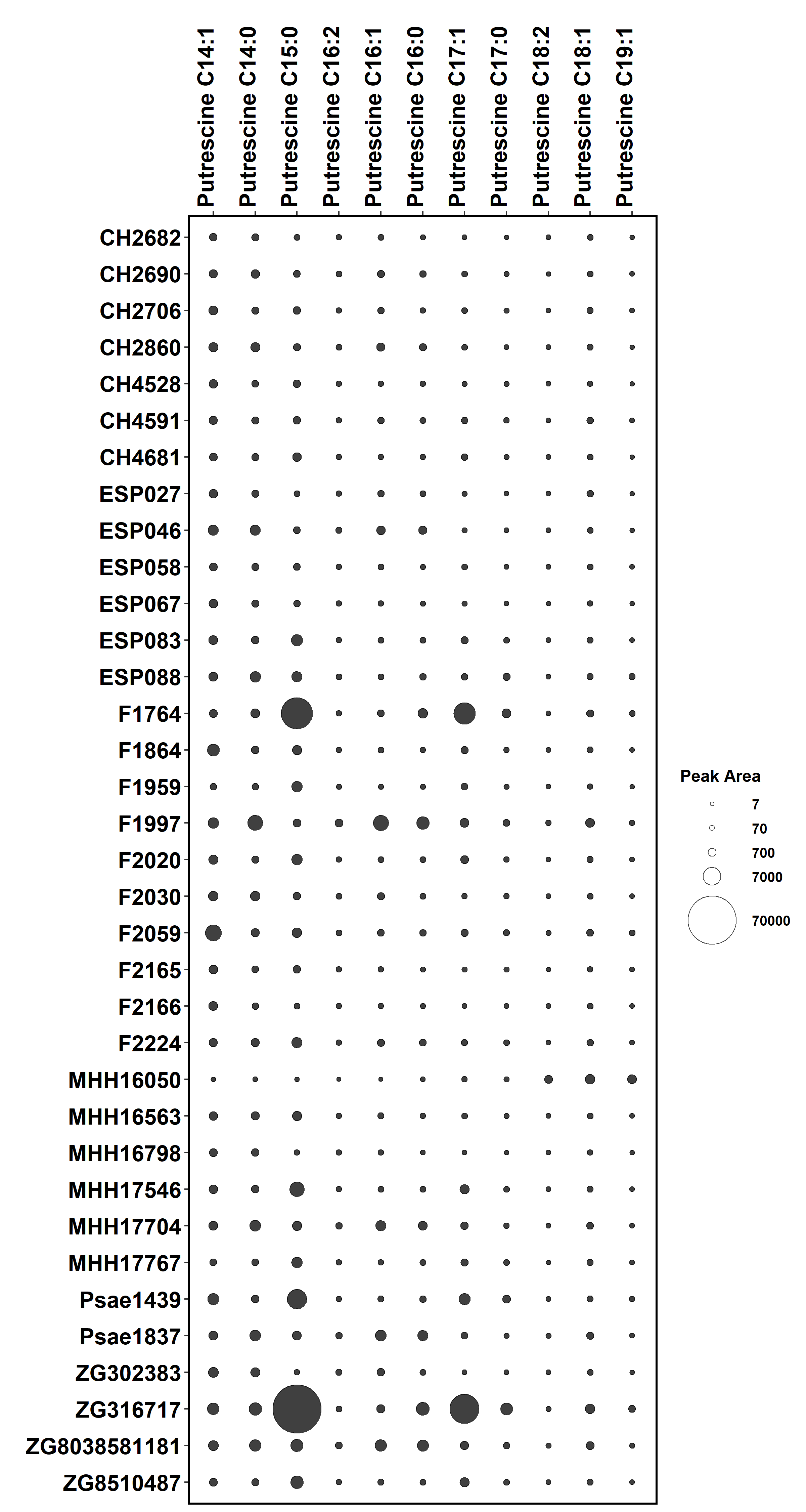

Figure S4. Bubble plot representing the quantitation (peak area) of eleven acyl putrescines from Dataset 2 (MSV000089869): publicly available LC-MS/MS data capturing the secondary metabolome of 35 P. aeruginosa isolates selected from the Helmholtz Centre for Infection Research biobank which were isolated from various sites of infection and cultured in LB under standard laboratory conditions (n = 1 biological replicate).

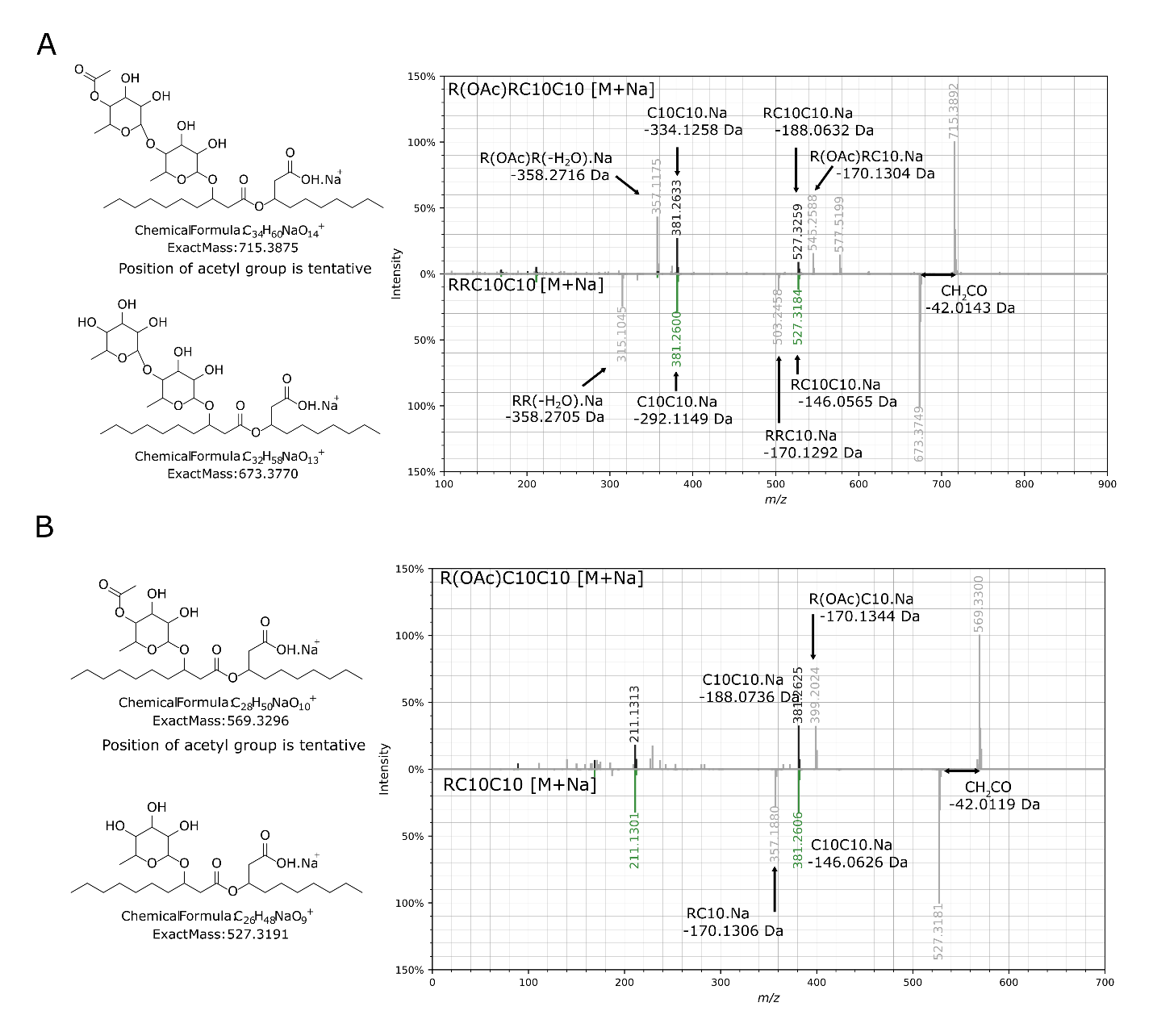

Figure S5. Annotation of the MS/MS spectra for the acetylated rhamnolipids (A) R(OAc)RC10C10 and (B) R(OAc)C10C10 compared to structurally characterized rhamnolipids RRC10C10 and RC10C10. Localization of acetyl group is putative.

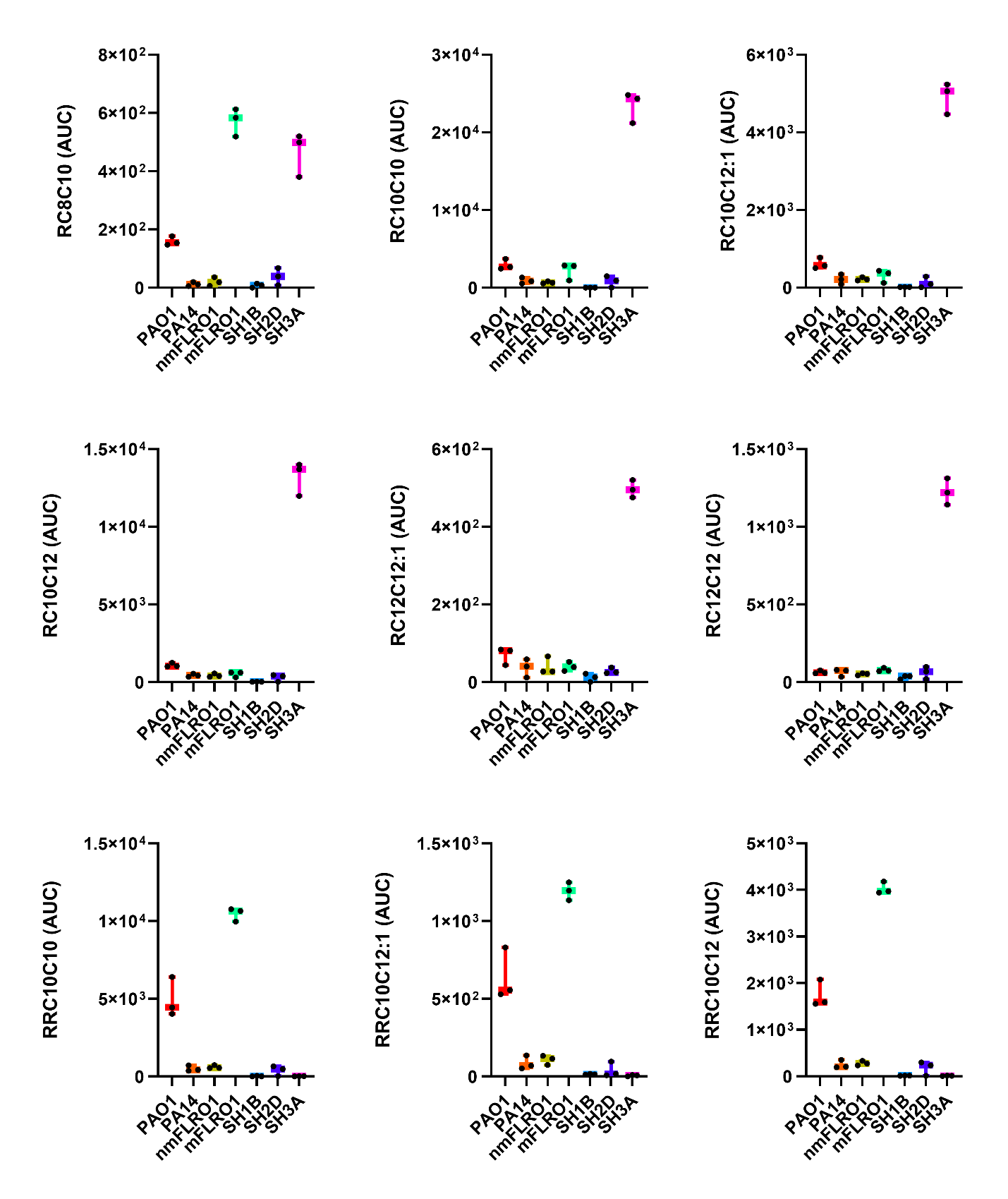

Figure S6. Differential levels of rhamnolipids produced by seven P. aeruginosa strains in SCFM2. Box plots represent the 25 to 75th percentiles, with a line at the median. Error bars indicate the minimum to maximum. Individual sample values shown (n = 3 biological replicates per strain).

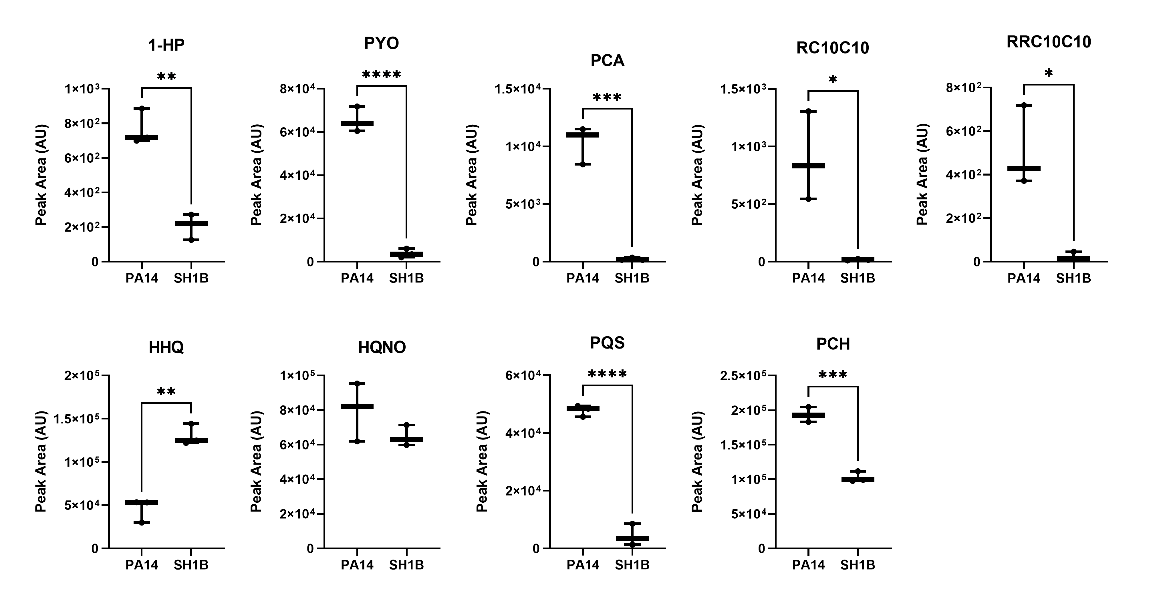

Figure S7. Differential levels of secondary metabolites produced by PA14 and SH1B in SCFM2. PCN was below the limit of quantitation. Box plots represent the 25 to 75th percentiles, with a line at the median. Error bars indicate the minimum to maximum. Individual sample values shown (n = 3 biological replicates per strain). Unpaired two-tailed T-tests. * p < 0.05; ** p < 0.01; *** p < 0.005; **** p < 0.001

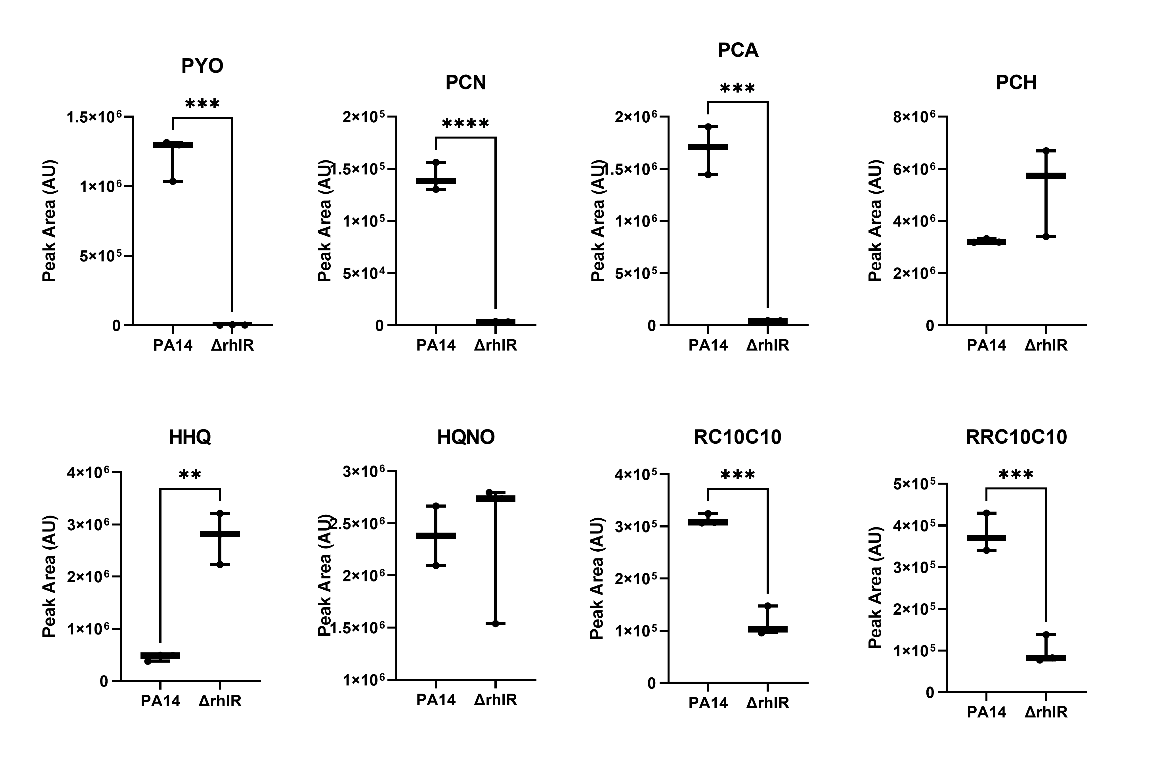

Figure S8. Differential levels of secondary metabolites produced by PA14 and ΔrhlR in LB (MSV000083500). 1‑HP and PQS were below the limit of quantitation. Box plots represent the 25 to 75th percentiles, with a line at the median. Error bars indicate the minimum to maximum. Individual sample values shown (n = 3 biological replicates per strain). Unpaired two-tailed T-tests. * p < 0.05; ** p < 0.01; *** p < 0.005; **** p < 0.001

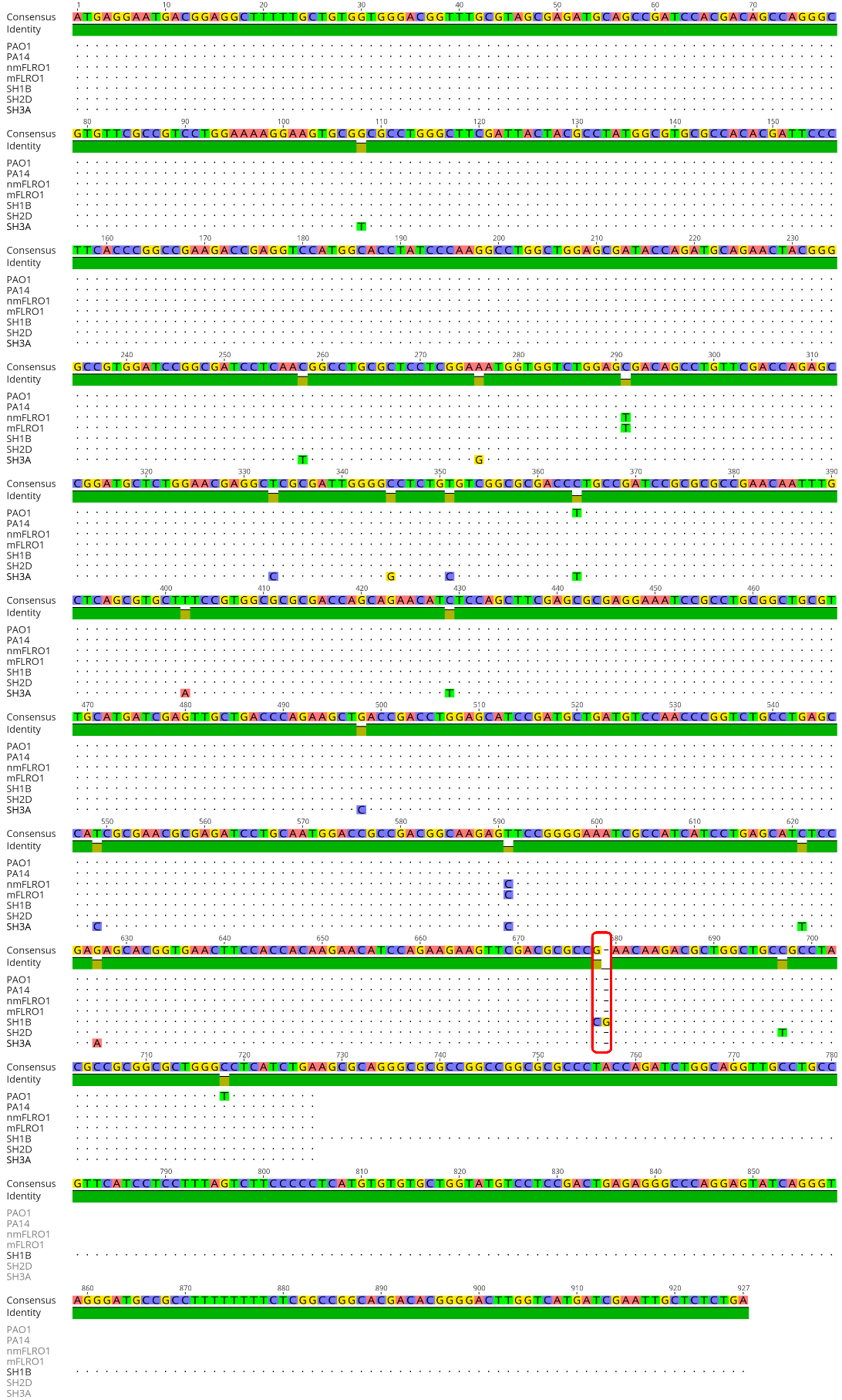

Figure S9. The rhlR gene sequences of PAO1, PA14, nmFLRO1, mFLRO1, SH1B, SH2D, and SH3A, with a red box highlighting single nucleotide insertion at position 678 in SH1B.

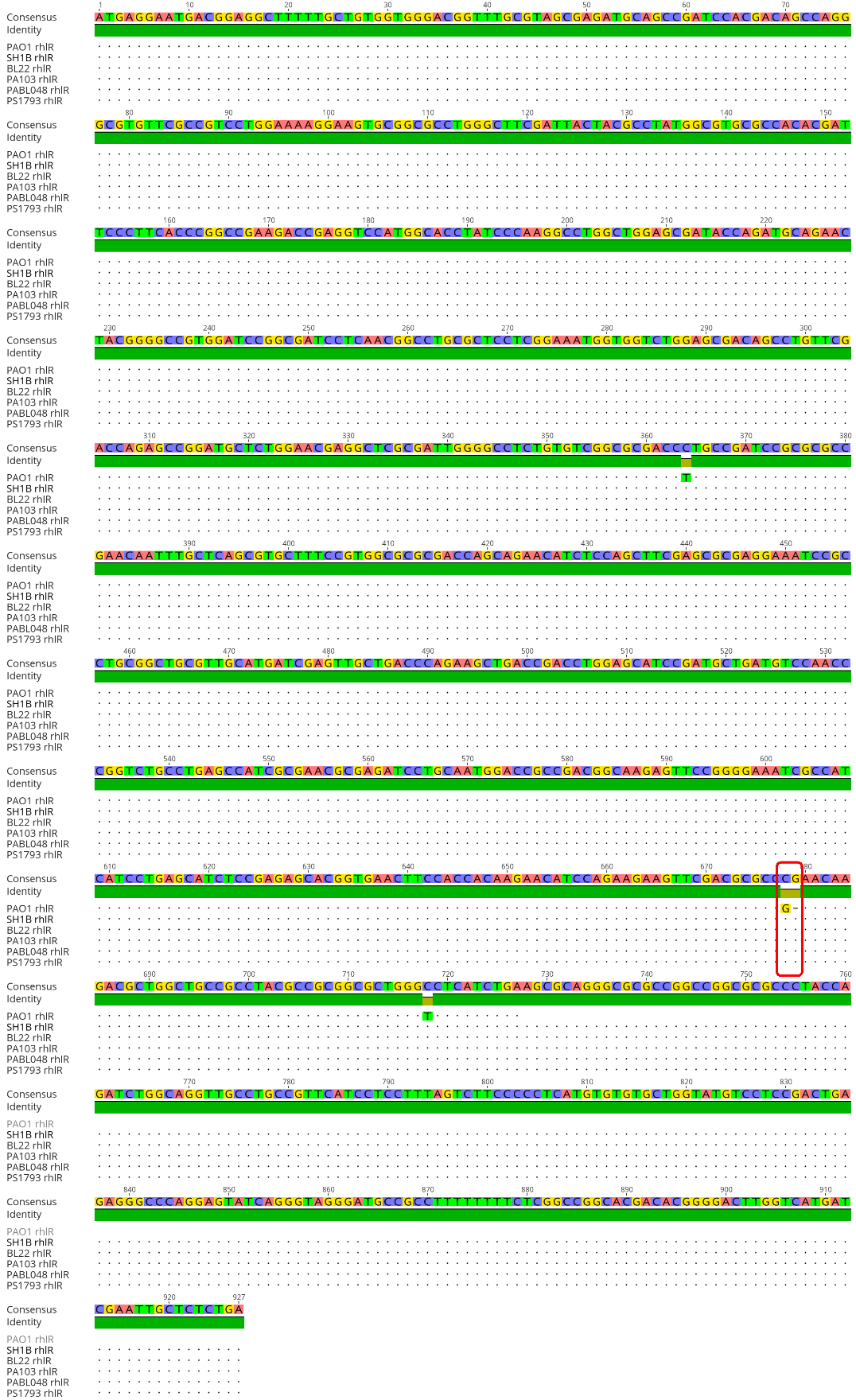

Figure S10. The rhlR gene sequences of PAO1, SH1B, BL22, PA103, PABL048, and PS1793, with a red box highlighting single nucleotide insertion at position 678 in the isolates compared to PAO1.

**Table S2. Secondary Metabolite Annotation**

| **Molecular Family** | **CMN Cluster ID** | **Annotation** | **Adduct** | **Measured m/z** | **RT** | **Calculated m/z** | **Mass Defect (ppm)** | **Annotation Level^#^** |
| --- | --- | --- | --- | --- | --- | --- | --- | --- |
| Phenazine | 462;463;465;466;467;468;469;480;488;512;541 | 1-HP | [M+H] | 197.0718 | 4.2 | 197.0709 | 4.57 | 1 |
| Phenazine | 828;829;831 | PYO | [M+H] | 211.0849 | 0.7 | 211.0866 | -8.05 | 1 |
| Phenazine | 828;829;831 | PYO | [M+H] | 211.0859 | 2.0 | 211.0866 | -3.32 | 1 |
| Phenazine | 1006 | PCN | [M+H] | 224.0822 | 4.0 | 224.0818 | 1.79 | 1 |
| Phenazine | 1018;1019;1020;1021;1022;1035 | PCA | [M+H] | 225.0659 | 4.5 | 225.0659 | 0 | 1 |
| Quinolone | 899;903;910 | C5-HQ | [M+H] | 216.1382 | 4.2 | 216.1383 | -0.46 | 3 |
| Quinolone | 975 | C4-QNO | [M+H] | 218.1180 | 3.8 | 218.1176 | 1.83 | 3 |
| Quinolone | 1236 | C6-HQ | [M+H] | 230.1536 | 4.6 | 230.1539 | -1.3 | 3 |
| Quinolone | 1390 | C5-QNO | [M+H] | 232.1332 | 4.4 | 232.1332 | 0 | 3 |
| Quinolone | 1529 | C7-HQ (HHQ) | [M+H] | 244.1694 | 5.2 | 244.1696 | -0.82 | 1 |
| Quinolone | 1571 | C6-QNO | [M+H] | 246.1488 | 4.8 | 246.1489 | -0.41 | 3 |
| Quinolone | 1830 | C8:1-HQ | [M+H] | 256.1697 | 5.7 | 256.1696 | 0.39 | 3 |
| Quinolone | 1849 | C7:1-QNO | [M+H] | 258.1488 | 5.1 | 258.1489 | -0.39 | 3 |
| Quinolone | 1849 | C8:1-HQ | [M+H] | 258.1851 | 5.7 | 258.1852 | -0.39 | 3 |
| Quinolone | 1861;1857;1864;1856;1858;1859;1862 | C7-QNO (HQNO) | [M+H] | 260.1642 | 5.3 | 260.1645 | -1.15 | 1 |
| Quinolone | 1861;1857;1864;1856;1858;1859;1862 | PQS | [M+H] | 260.1643 | 5.6 | 260.1645 | -0.77 | 1 |
| Quinolone | 1938;1937 | C9:1-HQ | [M+H] | 270.1851 | 6.1 | 270.1852 | -0.37 | 2 |
| Quinolone | 2133 | C8:1-QNO | [M+H] | 272.1643 | 5.5 | 272.1645 | -0.73 | 2 |
| Quinolone | 2138;2135 | C9-HQ (NHQ) | [M+H] | 272.2005 | 6.1 | 272.2009 | -1.47 | 2 |
| Quinolone | 2498 | C8-QNO | [M+H] | 274.1800 | 5.7 | 274.1802 | -0.73 | 2 |
| Quinolone | 2526 | PQS-OH | [M+H] | 276.1596 | 5.9 | 276.1594 | 0.72 | 3 |
| Quinolone | 2668;2670 | C10:1-HQ | [M+H] | 284.2011 | 6.1 | 284.2009 | 0.7 | 3 |
| Quinolone | 2668;2670 | C10:1-HQ | [M+H] | 284.2011 | 6.6 | 284.2009 | 0.7 | 3 |
| Quinolone | 2709;2710;2711;2712 | C9:1-QNO | [M+H] | 286.1799 | 5.9 | 286.1802 | -1.05 | 2 |
| Quinolone | 2709;2710;2711;2712 | C9:1-PQS | [M+H] | 286.1800 | 7.0 | 286.1802 | -0.7 | 3 |
| Quinolone | 2738;2739;2740;2741 | C9-PQS | [M+H] | 288.1956 | 6.5 | 288.1956 | 0 | 2 |
| Quinolone | 2738;2739;2740;2741 | C9-QNO (NQNO) | [M+H] | 288.1956 | 6.1 | 288.1956 | 0 | 3 |
| Quinolone | 2738;2739;2740;2741 | C9-HQ-OH | [M+H] | 288.1957 | 5.0 | 288.1956 | 0.35 | 3 |
| Quinolone | 2840 | C11:2-HQ | [M+H] | 296.2010 | 6.6 | 296.2009 | 0.34 | 3 |
| Quinolone | 2896 | C11:1-HQ | [M+H] | 298.2162 | 6.5 | 298.2165 | -1.01 | 2 |
| Quinolone | 2943 | C10:1-QNO | [M+H] | 300.1961 | 6.2 | 300.1958 | 1 | 2 |
| Quinolone | 2943 | C11-HQ | [M+H] | 300.2318 | 7.0 | 300.2322 | -1.33 | 2 |
| Quinolone | 3051;3053 | C9-QNO-OH | [M+H] | 304.1908 | 5.2 | 304.1907 | 0.33 | 3 |
| Quinolone | 3051;3053 | C9-PQS-OH | [M+H] | 304.1909 | 6.6 | 304.1907 | 0.66 | 3 |
| Quinolone | 3153;3176 | C11:2-QNO/C11:1-PQS | [M+H] | 312.1956 | 6.4 | 312.1958 | -0.64 | 3 |
| Quinolone | 3153;3176 | C12:1-HQ | [M+H] | 312.2320 | 6.9 | 312.2322 | -0.64 | 3 |
| Quinolone | 3207;3210 | C11:1-QNO | [M+H] | 314.2110 | 6.5 | 314.2115 | -1.59 | 3 |
| Quinolone | 3207;3210 | C11:1-PQS | [M+H] | 314.2114 | 6.8 | 314.2115 | -0.32 | 3 |
| Quinolone | 3230 | C11-QNO | [M+H] | 316.2268 | 7.0 | 316.2271 | -0.95 | 3 |
| Quinolone | 3283 | C13:2-HQ | [M+H] | 324.2316 | 7.3 | 324.2322 | -1.85 | 3 |
| Quinolone | 3321 | C13:1-HQ | [M+H] | 326.2475 | 7.3 | 326.2479 | -1.23 | 3 |
| Quinolone | 3409;3414 | C12:1-QNO/C12:1-PQS | [M+H] | 328.2273 | 6.9 | 328.2271 | 0.61 | 3 |
| Quinolone | 3517 | C13:2-QNO/C13:2-PQS | [M+H] | 340.2271 | 7.0 | 340.2271 | 0 | 3 |
| Quinolone | 3554 | C13:1-QNO | [M+H] | 342.2430 | 7.2 | 342.2428 | 0.58 | 3 |
| Quinolone | 3571 | C15:1-HQ | [M+H] | 354.2785 | 8.1 | 354.2791 | -1.69 | 3 |
| Quinolone | 3762 | C15-HQ | [M+H] | 356.2950 | 8.8 | 356.2948 | 0.56 | 3 |
| Quinolone | 4750 | C17:1-HQ | [M+H] | 382.3098 | 8.9 | 382.3104 | -1.57 | 3 |
| Siderophore | 3284 | PCH | [M+H] | 325.0673 | 4.6 | 325.0675 | -0.62 | 2 |
| Siderophore | 3284 | PCH | [M+H] | 325.0672 | 5.0 | 325.0675 | -0.92 | 2 |
| Rhamnolipid | 5930 | RC8C10 | [M+Na] | 499.2882 | 6.6 | 499.2878 | 0.8 | 1 |
| Rhamnolipid | 6052 | RC10C10 | [M+Na] | 527.3186 | 7.2 | 527.3191 | -0.95 | 1 |
| Rhamnolipid | 6269 | RC10C11 | [M+Na] | 541.3336 | 7.6 | 541.3347 | -2.03 | 1 |
| Rhamnolipid | 6806 | RC10C12:1 | [M+Na] | 553.3348 | 7.7 | 553.3347 | 0.18 | 1 |
| Rhamnolipid | 6812 | RC10C12 | [M+Na] | 555.3498 | 8.0 | 555.3504 | -1.08 | 1 |
| Rhamnolipid | 6878 | R(OAc)C10C10 | [M+Na] | 569.3300 | 7.8 | 569.3290 | 1.76 | 3 |
| Rhamnolipid | 6898 | RC12C12:1 | [M+Na] | 581.3648 | 8.5 | 581.3660 | -2.06 | 1 |
| Rhamnolipid | 6900 | RC12C12 | [M+Na] | 583.3820 | 8.8 | 583.3817 | 0.51 | 1 |
| Rhamnolipid | 8214 | RRC10C10 | [M+Na] | 673.3772 | 6.7 | 673.3770 | 0.3 | 1 |
| Rhamnolipid | 8367 | RRC10C12:1 | [M+Na] | 699.3922 | 7.2 | 699.3926 | -0.57 | 1 |
| Rhamnolipid | 8372 | RRC10C12 | [M+Na] | 701.4073 | 7.4 | 701.4083 | -1.43 | 1 |
| Rhamnolipid | 8377 | R(OAc)RC10C10 | [M+Na] | 715.3870 | 7.3 | 715.3875 | -0.7 | 3 |
| Rhamnolipid | 8395 | RRC12C12 | [M+Na] | 729.4410 | 8.2 | 729.4396 | 1.92 | 1 |
| Acyl Putrescine | 2862 | Putrescine C14:1 | [M+H] | 297.2898 | 5.3 | 297.2900 | -0.67 | 3 |
| Acyl Putrescine | 2935 | Putrescine C14:0 | [M+H] | 299.3061 | 5.7 | 299.3057 | 1.34 | 3 |
| Acyl Putrescine | 3206 | Putrescine C15:0 | [M+H] | 313.3217 | 5.8 | 313.3213 | 1.28 | 3 |
| Acyl Putrescine | 3282 | Putrescine C16:2 | [M+H] | 323.3060 | 5.7 | 323.3057 | 0.93 | 3 |
| Acyl Putrescine | 3314 | Putrescine C16:1 | [M+H] | 325.3216 | 5.8 | 325.3213 | 0.92 | 2 |
| Acyl Putrescine | 3388 | Putrescine C16:0 | [M+H] | 327.3373 | 6.2 | 327.3370 | 0.92 | 3 |
| Acyl Putrescine | 3546 | Putrescine C17:1 | [M+H] | 339.3374 | 6.0 | 339.3370 | 1.18 | 3 |
| Acyl Putrescine | 3661 | Putrescine C18:2 | [M+H] | 351.3376 | 6.2 | 351.3370 | 1.71 | 3 |
| Acyl Putrescine | 3705 | Putrescine C18:1 | [M+H] | 353.3529 | 6.3 | 353.3526 | 0.85 | 2 |
| Acyl Putrescine | 3945 | Putrescine C19:1 | [M+H] | 367.3690 | 6.4 | 367.3683 | 1.91 | 3 |
